# Leveraging Single-Cell RNA-Seq to Generate Robust Microglia Aging Clocks

**DOI:** 10.1101/2024.10.05.616811

**Authors:** Natalie Stanley, Luvna Dhawka, Sneha Jaikumar, Yu-Chen Huang, Anthony S Zannas

**Author notes:** These authors contributed equally to this work (Natalie Stanley, Luvna Dhawka).

## Abstract

‘Biological aging clocks’ - composite molecular markers thought to capture an individual’s biological age - have been traditionally developed through bulk-level analyses of mixed cells and tissues. However, recent evidence highlights the importance of gaining single-cell-level insights into the aging process. Microglia are key immune cells in the brain shown to adapt functionally in aging and disease. Recent studies have generated single-cell RNA sequencing (scRNA-seq) datasets that transcriptionally profile microglia during aging and development. Leveraging such datasets, we develop and compare computational approaches for generating transcriptome-wide summaries to establish robust microglia aging clocks. Our results reveal that unsupervised, frequency-based featurization approaches strike a balance in accuracy, interpretability, and computational efficiency. We further extrapolate and demonstrate applicability of such microglia clocks to readily available bulk RNA-seq data with environmental inputs. Single-cell-derived clocks can yield insights into the determinants of brain aging, ultimately promoting interventions that beneficially modulate health and disease trajectories.

## 1 Introduction

An exciting advancement over the recent years has been the development of ‘biological aging clocks’, composite molecular markers that are thought to capture the rate at which an individual ages biologically [1, 2]. Initially developed as predictors of chronological age [3, 4], the subsequently developed clocks were further shown to predict diverse aging-related disease and mortality outcomes [5–7], supporting their promise as disease biomarkers. To date, aging clocks have been generated by various molecular platforms - e.g., transcriptomics, epigenomics, proteomics - largely applied in data combined from many tissues and organs or in heterogeneous mixtures of cells from single tissues (i.e., at the bulk level) [3, 8]. While such bulk-level analyses are essential for biomarker discovery, molecular changes and aging rates are known to vary across different tissues and cell types [9–11]. Moreover, disease-related alterations in aging clocks and their potential underlying pathways have been reported to occur in a tissue- and cell-type-specific manner [12–14], highlighting the importance of gaining cell-level insights into biological aging.

Microglia are key immune cells in the brain [15, 16] and have been critically implicated in the neuroinflammation associated with aging [17, 18], adaptations to stress [19, 20], neurodegenerative states [21–23], and diverse neuropsychiatric diseases [24, 25]. In these processes, microglia can undergo dynamic transcriptional alterations that are relevant to brain function and disease pathogenesis [19, 23, 26]. Recent seminal single-cell RNA sequencing (scRNA-seq) studies further suggest that such transcriptional alterations involve specific microglia subtypes and key genetic programs that can be leveraged by machine learning algorithms to develop microglia aging clocks [27– 30]. Importantly, functional alterations in microglia subtypes are also associated with distinct aging-related brain phenotypes, including neural stem cell proliferation [28], neurodegeneration [31], and Alzheimer’s disease [30]. Uncovering the transcriptional dynamics of microglia at the single-cell and cell-type levels may thus yield unique insights into the determinants of brain aging and disease.

With single-cell technologies offering an unprecedented level of resolution in characterizing functional states of individual cells, quality computational methods are required to translate such information into machine learning models of sample-level phenotype. Here, *featurization* denotes the process of translating multiple prominent gene expression patterns and relative cell-type abundances from single-cell datasets into succinct summaries that can be input to machine learning models of aging. Practically, each sample profiled with a single-cell technology produces a large matrix of many features measured per individual cell, which must be translated into a sample-level feature vector to ultimately train machine learning models of age. Computational featurizations of single-cell data have been explored substantially for translating abundances and functional states of immune-cell types assayed with flow and mass cytometry [32–39], but these approaches are not as well explored for high-dimensional scRNA-seq data. While supervised featurization methods can be trained to learn per-sample representations, based on external information such as age [36, 37], here we examine the capacity of unsupervised feature engineering strategies, such as computing cell-type frequencies [32] or pseudobulk-level features [40] to be used to train single-cell microglia based models of age. Such unsupervised approaches are well suited for datasets with small sample sizes and for uncovering meaningful patterns that are biologically interpretable, such that they can illuminate the cell types associated with particular aging trajectories. While Buckley *et al*. pioneered the use of a pseudobulk-based approach to compute per-sample featurizations across diverse cell-types in the brain, it has not been adequately explored how the myriad of featurization approaches can translate cellular heterogeneity patterns into models of age.

In this study, we leveraged publicly available scRNA-seq datasets profiling multiple microglia samples across the lifespan (Fig. 1**a**), to pursue three main objectives. First, we determined the extent to which frequencies of particular microglia subtypes and their gene expression patterns change dynamically during aging and development (Fig. 1**b**). Using the same datasets, we also quantified how four methodologically diverse, unsupervised featurization approaches of microglia transcriptome-wide signatures perform as age classifiers and aging clocks (Fig. 1**c**). Lastly, we assessed whether the newly constructed microglia clocks and their conserved genetic programs are applicable to bulk RNA-seq data with environmental inputs (Fig. 1**d**). Overall our findings suggest that single-cell-derived and cell-type-specific microglia clocks can yield unique, biologically relevant insights into brain aging.

**Fig. 1.**
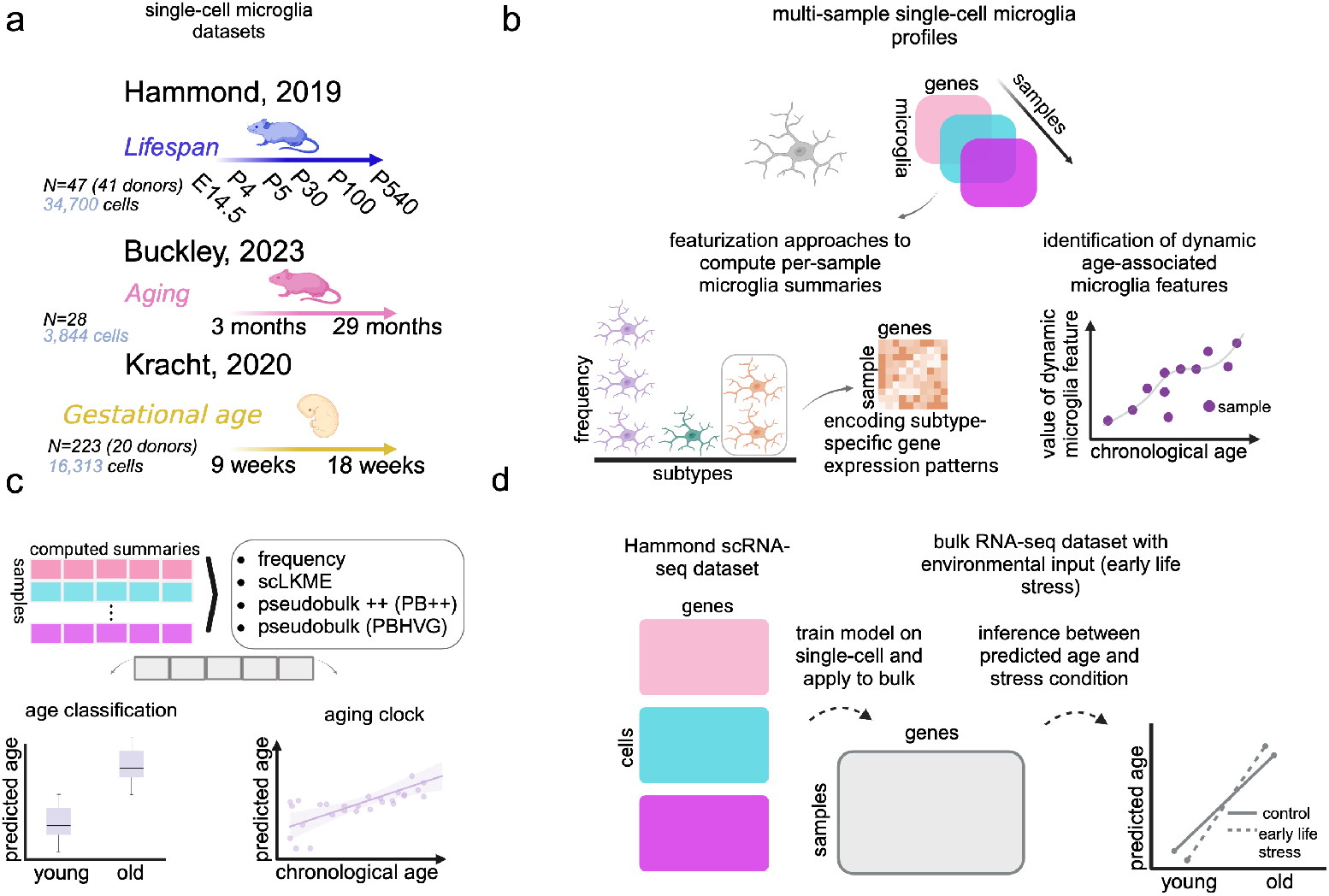
Overview. **a** We used three scRNA-seq datasets profiling microglia during aging and development to build microglia aging clocks. The Hammond dataset has six discrete time points spanning between embryonic stage (E14.5) and old age (P540). The Buckley and Kracht datasets have continuously sampled ages between 3 and 29 months and 9 and 18 gestational weeks, respectively. **b** To ultimately build aging clocks, or machine learning models of age, we computed computational summaries or *featurizations* for each sample based on frequencies and genetic programs of microglia subtypes. **c** Four approaches for featurizing or summarizing transcriptomic patterns of microglia in profiled samples were applied and include frequency, single-cell landmark kernel mean embedding (scLKME), pseudobulk++ (PB++), and classical pseudobulk (PBHVG). The various featurization methods generate succinct features that can be used as input to machine learning models of age. **d** A model for age was trained on the Hammond single-cell data and applied to an independent bulk RNA-seq dataset with an additional environmental input (exposure to early life stress).

## 2 Results

To understand the trade-offs in accuracy, interpretability, and efficiency of different featurization approaches, we leveraged three multi-sample scRNA-seq datasets that profile microglia transcriptionally during lifespan [27], aging [28], and fetal development [41] (Fig. 1**a**). Key microglia subtypes and their genetic programs identified under the different featurization approaches were leveraged to generate signatures of microglia in aging, and to compare such signatures across multiple datasets. To assess whether the findings uncovered through single-cell analysis can be generalized at the bulk level, the prominent genes identified by scRNA-seq were validated and used to train an aging clock in a bulk RNA-seq dataset of young (P9) and old (P200) mice generated by Reemst *et al*. [42].

### 2.1 Single-Cell RNA-Sequencing Datasets of Microglia During Aging and Development

Here, we provide an overview of the three scRNA-seq datasets (Fig. 1**a**) used to construct the microglia aging clocks with the four unique featurization approaches (Fig. 1**b**).

#### Hammond Mouse Lifespan

The Hammond dataset profiles mice throughout the lifespan [27], with samples collected at embryonic day 14.5 (E14.5), and postnatal days 4 (P4), 5 (P5), 30 (P30), 100 (P100), and 540 (P540). Our analysis included a total of 34,700 microglia isolated by fluorescence-activated cell sorting (FACS) in *N* = 47 total mouse samples from 41 unique donors. Note that five of the donors had multiple samples due to additional experimental perturbations used to induce a demyelinating injury.

#### Buckley Mouse Aging

The Buckley mouse aging dataset [28] is comprised of *N* = 28 mice sampled at different ages between 3.3 months and 29 months. Our analysis included a total of 3,844 cells. In the original study, cells extracted from the subventricular zone (SVZ) neurogenic region were profiled with scRNA-seq, but we selected microglia according to marker genes indicated in the original study, including, *C1qb, C1qb, Cst3, C1qc, Ctss, Hexb, Fcer1g, Trem2*, and *Tyrobp*.

#### Kracht Fetal Development

The Kracht fetal development dataset [41] profiled microglia that were isolated by FACS in post-mortem brains from 20 aborted human fetuses, obtained at timepoints sampled at different ages between 9 and 18 weeks of gestational age. Each donor contributed between one and four samples, resulting in *N* = 223 total samples. Our analysis included a total of 16,313 cells.

### 2.2 Microglia subtype frequencies vary with age

All three datasets exhibited microglia heterogeneity based on Leiden clustering that identified diverse microglia subtypes with characteristic genetic programs (Supplementary Figure 1). As a first approach to link cell heterogeneity to age, we examined how the frequencies of identified microglia subtypes varied over the aging trajectory in each dataset. To do so, we computed frequency features as the fraction of each sample’s cells assigned to each cluster according to previously described methods [32]. To identify the microglia subtypes that most strongly associate with age, subtype frequencies were then used to train and test a Random Forest classifier for age group over 200 trials (see Methods), and mean Gini [43] scores were computed as a metric of frequency feature importance (Fig 2). Fig. 2 **left** shows cells from the Hammond, Buckley, and Kracht datasets (**a-c**, respectively) and colored by the importance of the frequency of the cluster to which they belonged. In each dataset, we further highlighted the trajectories of the top five clusters with predictive and dynamic frequencies across the age spectrum (Fig. 2 **middle**). In the Hammond dataset, clusters 4 and 11 exhibited similar patterns, having high frequencies at embryonic stage (E14.5), peaking at P4, and then attenuating at older ages. Cluster 4 showed high expression of *Ftl1, Apoe*, and *Ctsb* (Fig. 2**a**, **right**, Supplementary Figure 8), whereas cluster 11 was marked prominently by *Stmn1, Tubb5*, and *Tuba1a*. Cluster 13 increased in frequency with age and was marked by *Malat1, Apoe*, and *Ifitm3*. In the Buckley dataset, clusters 4 and 10 exhibited very similar gene expression patterns with frequencies that significantly increased between adult and old age. These were marked by expression of *Fth1, Ftl1* and *Ctsb* (Fig. 2**b**, **right**, Supplementary Figure 9). Clusters 3 and 8 also had distinct gene expression patterns with frequencies that were highest at young and adult ages but decreased in old age. These clusters were prominently marked by transcription factors *Jun* and *Junb*, as well as by *Malat1*. Finally, in the Kracht dataset, clusters 5, 4, and 9 increased in frequency between the first and second trimesters and were marked by expression of *Cx3cr1, Ftl*, and *Csf1r* (Fig. 2**c**, **right**, Supplementary Figure 10), whereas clusters 2 and 8 decreased in frequency with age and both expressed *Malat1, Ftl*, and *Spp1*. Together these analyses indicate dynamic changes in microglia subtypes and genetic programs across aging and developmental stages.

**Fig. 2.**
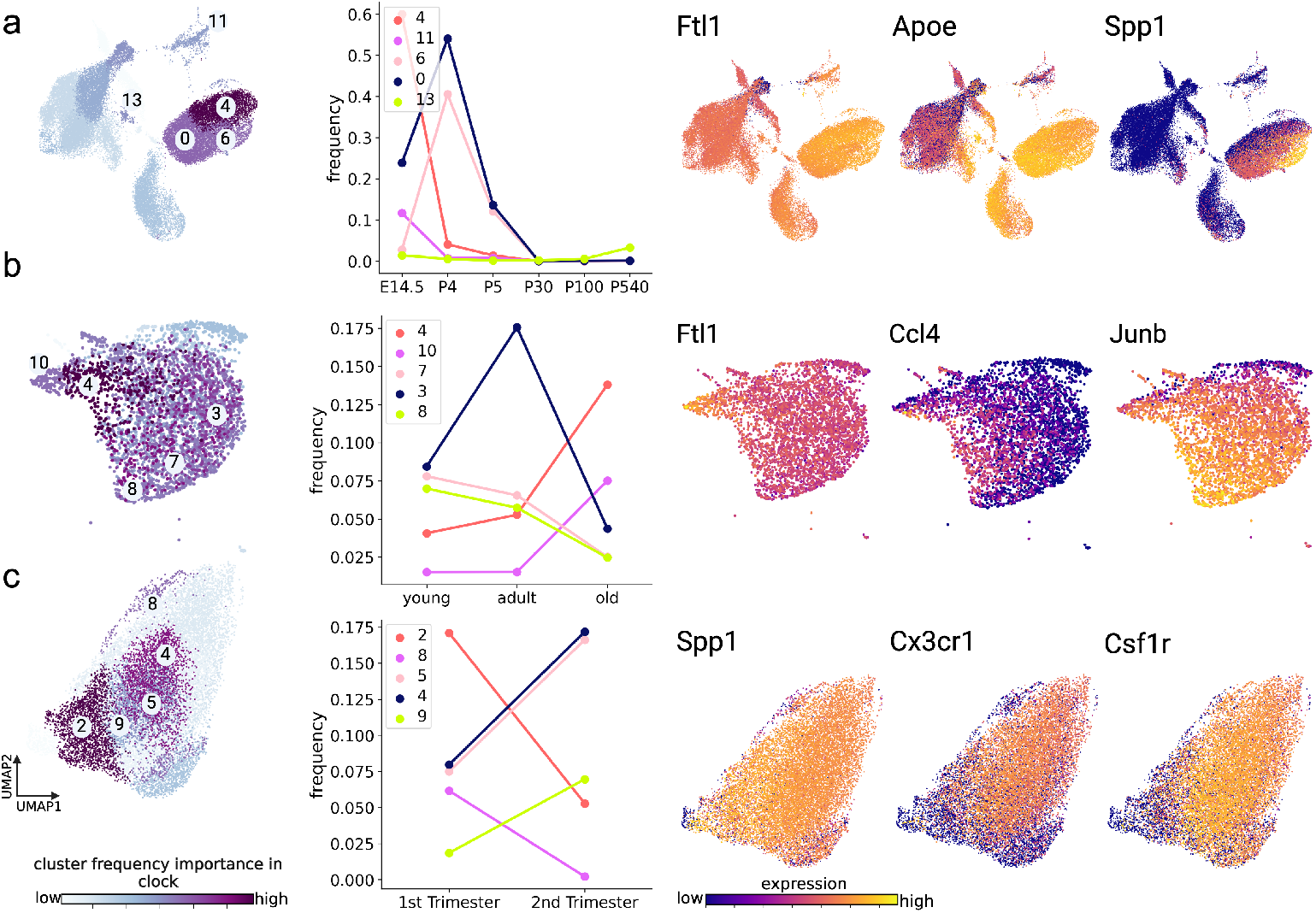
Frequencies of microglia subtypes change dynamically during aging and development. Age-related changes in microglia subtype frequencies were observed across the Hammond (**a**), Buckley (**b**), and Kracht (**c**) datasets. **Left** panels show two-dimensional UMAP projections of cells in each dataset. Cells are colored by the usefulness of their respective cluster’s frequency in predicting age (light to dark indicates less to more useful), according to cluster frequency Gini score computed for a random forest classifier trained to predict discrete age groups. Discrete age groups were defined for all datasets (further details in Methods). Five representative clusters with critical dynamic changes in frequency useful for age-classification are depicted and numbered in each UMAP plot. **Middle** panels plot the trajectory of mean frequencies of the top five age-predictive clusters in each dataset as a function of age group. **Right** panels show two-dimensional UMAP projections of cells in each dataset colored by the expression (dark to light indicates low to high expression) of key genes that were differentially expressed in at least one of the five prioritized clusters.

### 2.3 Featurization Methods Impact the Accuracy of Microglia Aging Markers

With evidence that microglia subtypes change over the aging continuum, we next explored a range of strategies to generate aging markers derived from scRNA-seq data. While clocks have historically been developed by modeling age as a continuous variable, we evaluated the usefulness of each featurization approach in both classifying age groups (i.e., discrete age variable) and predicting age (i.e., continuous age variable), through classification and regression models, respectively. More specifically, we examined how four methodologically distinct featurization techniques can be applied to translate a multi-sample single-cell dataset into a succinct vector representation or transcriptome-wide summary that can be used as input to a machine learning model of age (Fig. 1**c**). We provide a description of each featurization algorithm here and further technical details in the Methods section.

#### Frequency

Frequency-based featurization [32, 33] involves defining a common set of microglia subtypes within a sample through unsupervised clustering of all cells across samples within a dataset. A feature representation is computed for each sample by counting the fraction of its cells for each subtype.

#### scLKME

scLKME [39] is an unsupervised feature learning approach, which encodes the complexity of the single-cell landscape into a mathematically abstract vector representation that can be used as input to classification models. scLKME performs landmark-based kernel mean embedding, where a set of *landmarks* or generally prototypical cells are chosen across all samples. Similarity patterns of each cell in each sample are encoded based on overall similarity to the landmark cells, as evaluated through a kernel evaluation [35].

#### Pseudobulk based on highly variable genes (PBHVG)

Pseudobulk representations of single-cell data are the most common approach for summarizing broad gene expression patterns in single-cell data to create a representation similar to bulk data. While there are a range of techniques for creating pseudobulk profiles, outlined comprehensively in Ref. [40], here we computed the sum of each gene’s expression in each cluster. Because considering the expression of all measured genes would create a feature space with a prohibitively high dimensionality, we considered only the expression of the 3,000 most highly variable genes (HVG) in each cluster, which is a common pre-processing step in many single-cell pipelines [44]). This also implies that there are 3,000 generated pseudobulk features per cell cluster.

#### Pseudobulk++ (PB++)

In contrast to a traditional pseudobulk approach [40], which computes aggregate expression measurements for all genes across all cells in each cluster, the pseudobulk (PB) ++ approach computes a single value for each cluster, which reflects the general frequency and alignment with key gene expression programs of cells in the sample. Specifically, PB++ variants integrate frequency information (i.e., the number of cells across major cell-populations) with the extent to which key genes are expressed in each cluster. In the PB++25 and PB++50 variants, we use the top 25 and 50 differentially expressed genes across clusters, respectively, to imbue the feature representations with gene expression information. This newly proposed method aims to parsimoniously integrate signals originating from cell-type frequencies and key genetic programs.

#### 2.3.1 Microglia Age Classifiers

To assess classification performance across featurization approaches, ages in each dataset were binned into discrete categories, based on age group (Hammond), age distribution (Buckley), or trimester of pregnancy (Kracht). Histograms of age distributions are shown in Supplementary Figure 6. Note that the Hammond dataset already had discrete age categories (embryonic stage E14.5, P4, P5, P30, P100, and P540). The Buckley dataset was discretized into young (*<* 4 months), adult (4 months to less than 14 months), and old (≥ 14 months and above). The Kracht dataset was binned by trimester, such that 1st and 2nd trimesters contained samples of ages 9-12 and 13-18 weeks, respectively. Classification experiments involved 200 trials of random train/test splits, where 80% of donors and their respective samples were used for training and the remaining 20% for testing model accuracy (Acc), defined as the fraction of samples with a correctly predicted age group. Fig. 3 shows the results of the classification experiments across featurization approaches and datasets. In the Hammond dataset, which has the largest number of samples and age groups, the top performing methods were frequency and PB++, resulting in mean accuracies of Acc = 0.840 and Acc = 0.813, respectively. This suggests that larger datasets are sufficiently well-powered to use these simple, low-dimensional featurization approaches. In the Buckley dataset, which has the smallest number of samples, the top performing method was scLKME (mean Acc = 0.752), followed by PB++50 (mean Acc = 0.693). This suggests that features based on similarity patterns with landmark or anchor cells contained enough signal to accommodate the relatively small sample size. The Kracht dataset achieved strong performance with frequency features (mean Acc = 0.925), followed by PBHVG (mean Acc = 0.918). In each of the three datasets, either frequency or a PB++ variant achieved the top two highest classification accuracies. Such methods are the least high-dimensional, with the number of features corresponding to the number of clusters or inferred microglia sub-types. Therefore, our results suggest robust classification performance by simple and computationally efficient approaches, such as frequency or PB++ variants across aging and development.

**Fig. 3.**
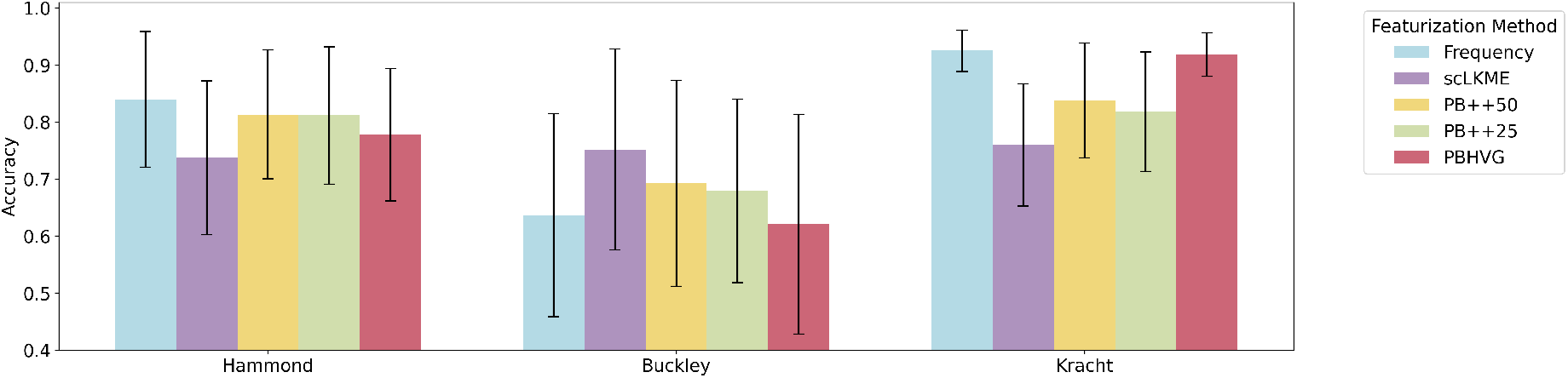
Accuracy of microglia aging classifiers across featurization approaches. Discrete age groups were created in each dataset to formulate a discrete age classification problem, using the various featurizations, including frequency, scLKME, Pseudobulk++50 (PB++50), Pseudobulk++25 (PB++25), and pseudobulk with highly variable genes (PBHVG) as input. Classification experiments were performed using random forest classifiers and repeated 200 times with randomized train/test splits of donors. Barplots and error bars show the mean and standard deviation in classification accuracy obtained over the 200 trials. Details on discrete age categories specified for each dataset are provided in Methods.

#### 2.3.2 Microglia Aging Clocks

To generate microglia aging clocks (i.e., predictors of chronological age), we employed a leave-one-out cross-validation approach that used all but one donor and their respective samples to train a Lasso Regression model on a given set of features. We then evaluated the predicted age of the held-out sample or donor and plotted them against their chronological age in Fig. 4. In the Hammond dataset (Fig. 4 **top**), PB++50 showed the highest correlation between chronological and predicted ages (*r* = 0.90, *p* = 3.04 *×* 10^−18^), followed by scLKME (*r* = 0.88, *p <* 6.87 *×* 10^−16^). In the Buckley dataset (Fig. 4 **middle**), frequency showed the highest concordance between chronological and predicted age (*r* = 0.82, *p* = 9.3 *×* 10^−8^), with scLKME as the second top-performing method (*r* = 0.80, *p* = 3 *×* 10^−7^), suggesting that the additional information content included through a higher-dimensional feature space was only marginally helpful.

**Fig. 4.**
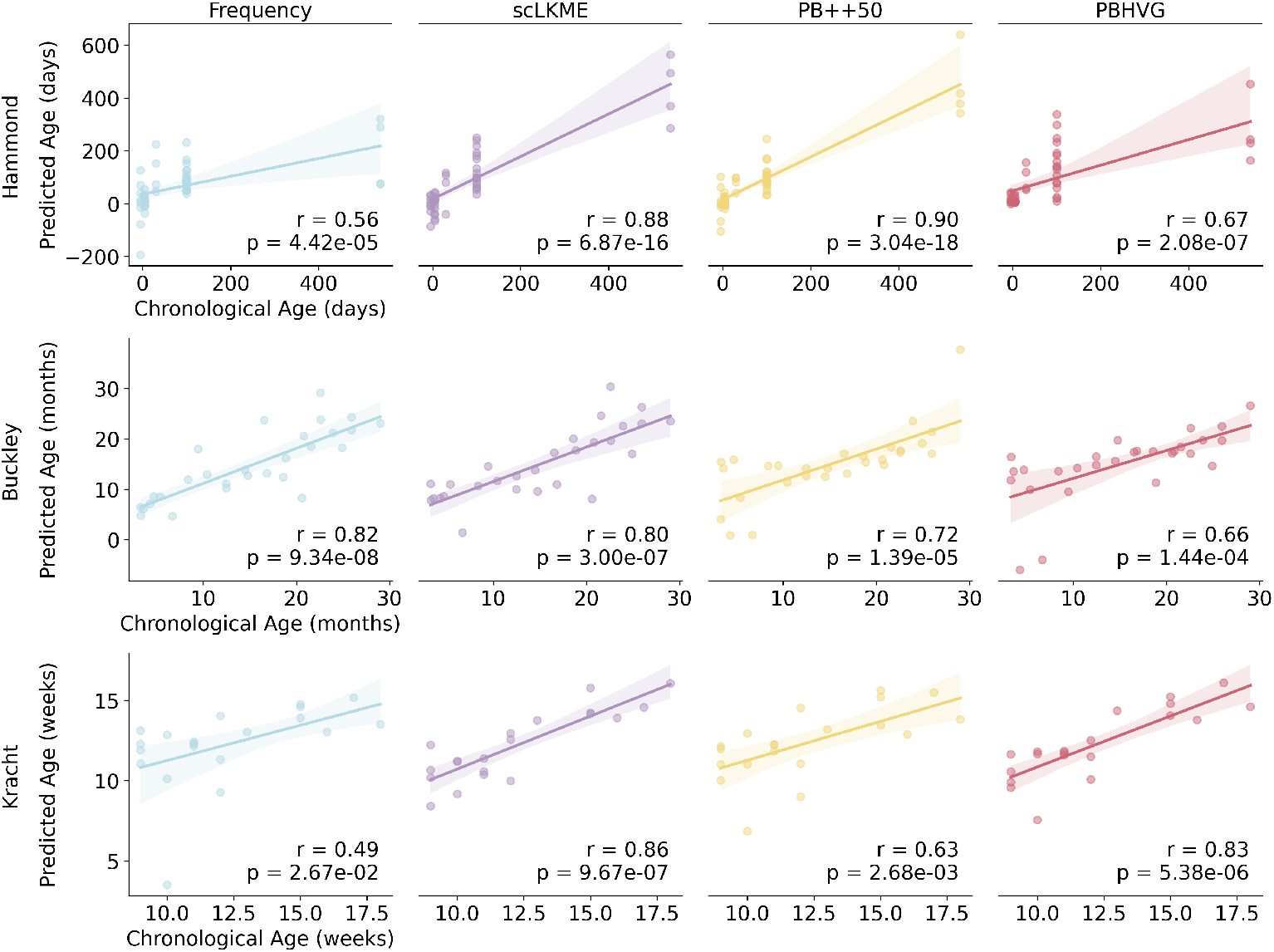
Performance of microglia aging clocks across featurization approaches. Age was used as a continuous variable to train Lasso regression models under frequency, scLKME, Pseudobulk++50 (PB++50), and pseudobulk with highly variable genes (PBHVG) featurization approaches, using leave-one-out cross validation (LOOCV) in the Hammond, Buckley, and Kracht datasets. Scatterplots show the predicted versus true chronological age for each sample obtained in the LOOCV trial for which it was the test sample. Note that the Kracht dataset has multiple samples (replicates) per donor, so each point depicts the mean predicted age of all samples for each donor from the LOOCV iteration in which it was a test sample. Correlation coefficients between chronological and predicted ages are quantified with Pearson correlation (*r*).

In the Kracht dataset, the regression approach was more challenging, given the limited age range (Fig. 4 **bottom**). scLKME achieved the highest accuracy (*r* = 0.86, *p* = 9.67 *×* 10^−7^), followed by PBHVG (*r* = 0.83, *p* = 5.38 *×* 10^−6^). Overall, the results suggest that for datasets covering a more extended age range (like Hammond and Buckley), scLKME is the most robust choice, but lower-dimensional methods such as PB++50 and frequency can also achieve robust prediction accuracies. In datasets with a more limited age range (like Kracht), higher-dimensional approaches such as PBHVG and scLKME may be necessary to maximize the signal to noise ratio necessary for robust prediction. It is also worth noting that frequency features showed the weakest performance in the Hammond and Kracht datasets, despite excelling in age classification (Fig. 3), suggesting these features may lack adequate transcriptional signal for the regression task. This is especially true in the Kracht dataset, where we hypothesize that the limited age range is not discernible through frequency features alone.

### 2.4 Computational Requirements of Featurization Approaches

We next compared the computational requirements in terms of both run-time and memory across the featurization approaches (with run-time and memory requirements shown in Fig. 5 **top** and **bottom**, respectively). The variation in run-time and in the resources required for age classification and prediction was primarily driven by the number of features produced under each approach (Table 1). Fig. 5 shows mean run-time (**top**) and memory requirements (**bottom**). The analogous results for using Lasso regression to build a clock from each featurization approach are shown in Supplementary Figure 7.

**Table 1.** The number of features generated under each combination of dataset and featurization approach.

|  | **Hammond** | **Buckley** | **Kracht** |
| --- | --- | --- | --- |
| **Frequency** | 17 | 14 | 13 |
| **scLKME** | 1500 | 1500 | 1500 |
| **PB++25 & PB++50** | 17 | 14 | 13 |
| **Pseudobulk (PBHVG)** | 51,000 | 42,000 | 21,905 |

**Fig. 5.**
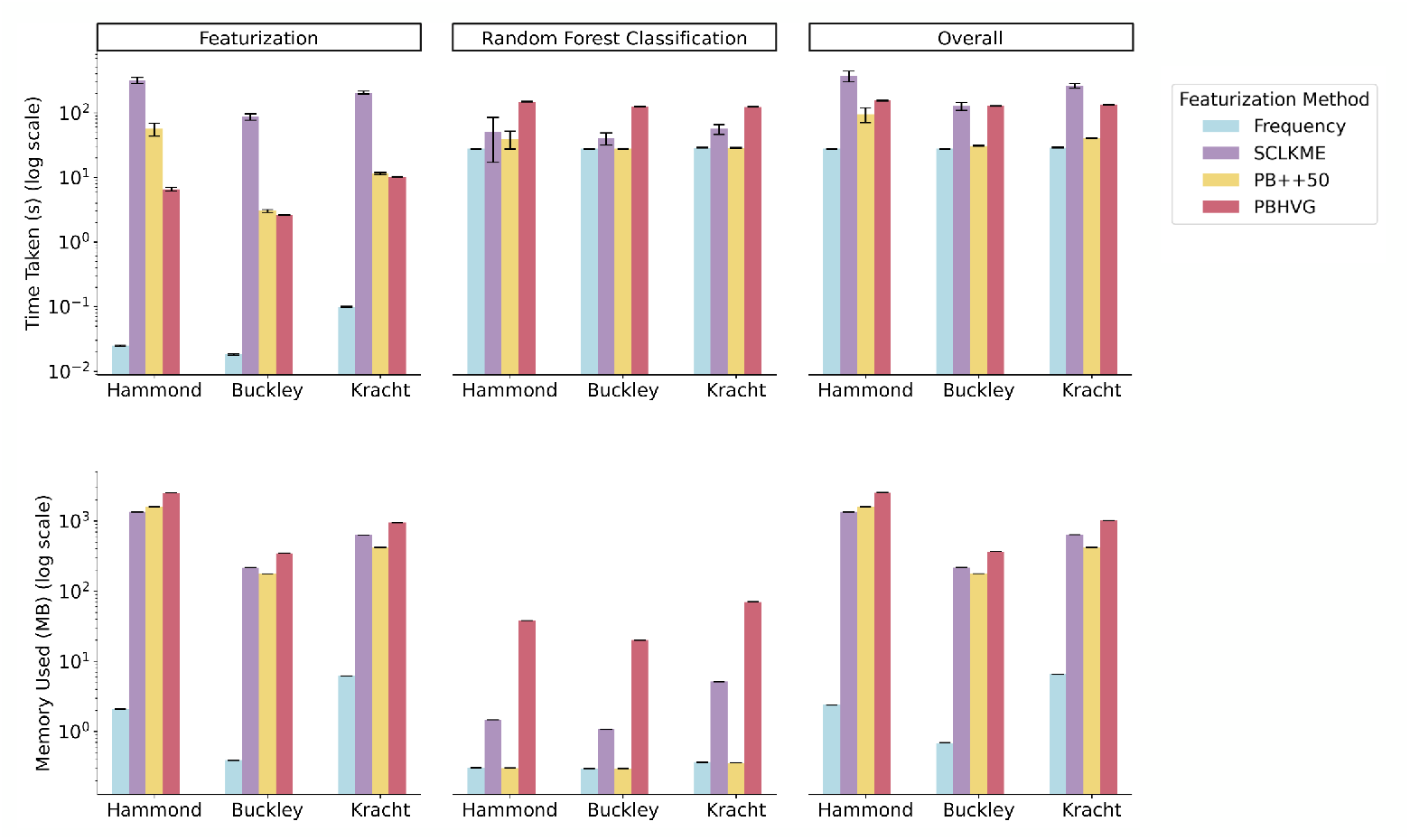
Run-time and memory requirements of featurization approaches. Run-time (**top**) and memory (**bottom**) required for each featurization approach, including frequency, scLKME, pseudobulk++50 (PB++50), and pseudobulk with highly variable genes (PBHVG) were evaluated across datasets for featurization only (**left**), age classification based on a Random Forest classifier (**middle**), and the overall process of featurization and classification (**right**). Results were obtained by repeating the featurization and model training over 30 trials and reporting the mean. Error bars show standard deviation around the mean. Results were obtained on a machine with Intel(R) Xeon(R) Gold 6226R CPU @ 2.90GHz.

**Fig. 6.**
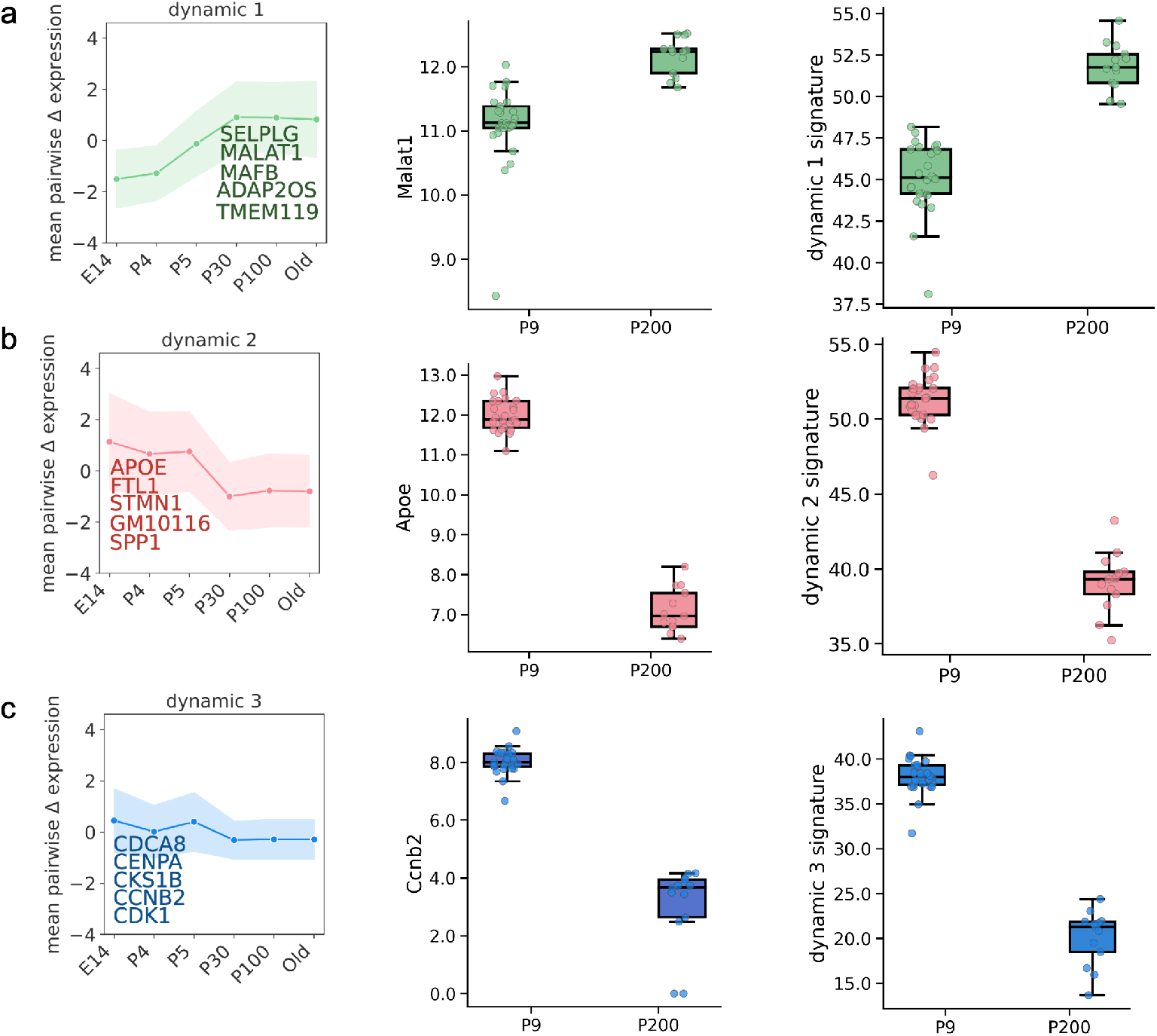
Modules of genes with characteristic dynamic changes in expression during aging are conserved across datasets. DELVE was used to identify modules of genes with common patterns of increase (**a**) or decrease (**b**,**c**) across the aging continuum in the Hammond dataset and applied to an independent bulk RNA-seq dataset. The top five genes for each module are highlighted. The **left** panel shows the average expression pattern across the mouse lifespan for five representative genes in each particular module. The **middle** panels show the distribution of expression patterns for a prominent, representative gene (*Malat1, Apoe*, and *Ccnb2*) in dynamic modules 1, 2, and 3, respectively in P9 and P200 mice. The **right** panels show distributions of aggregate (e.g. summed) gene expressions of the indicated top five genes in each module in P9 and P200 mice.

**Fig. 7.**
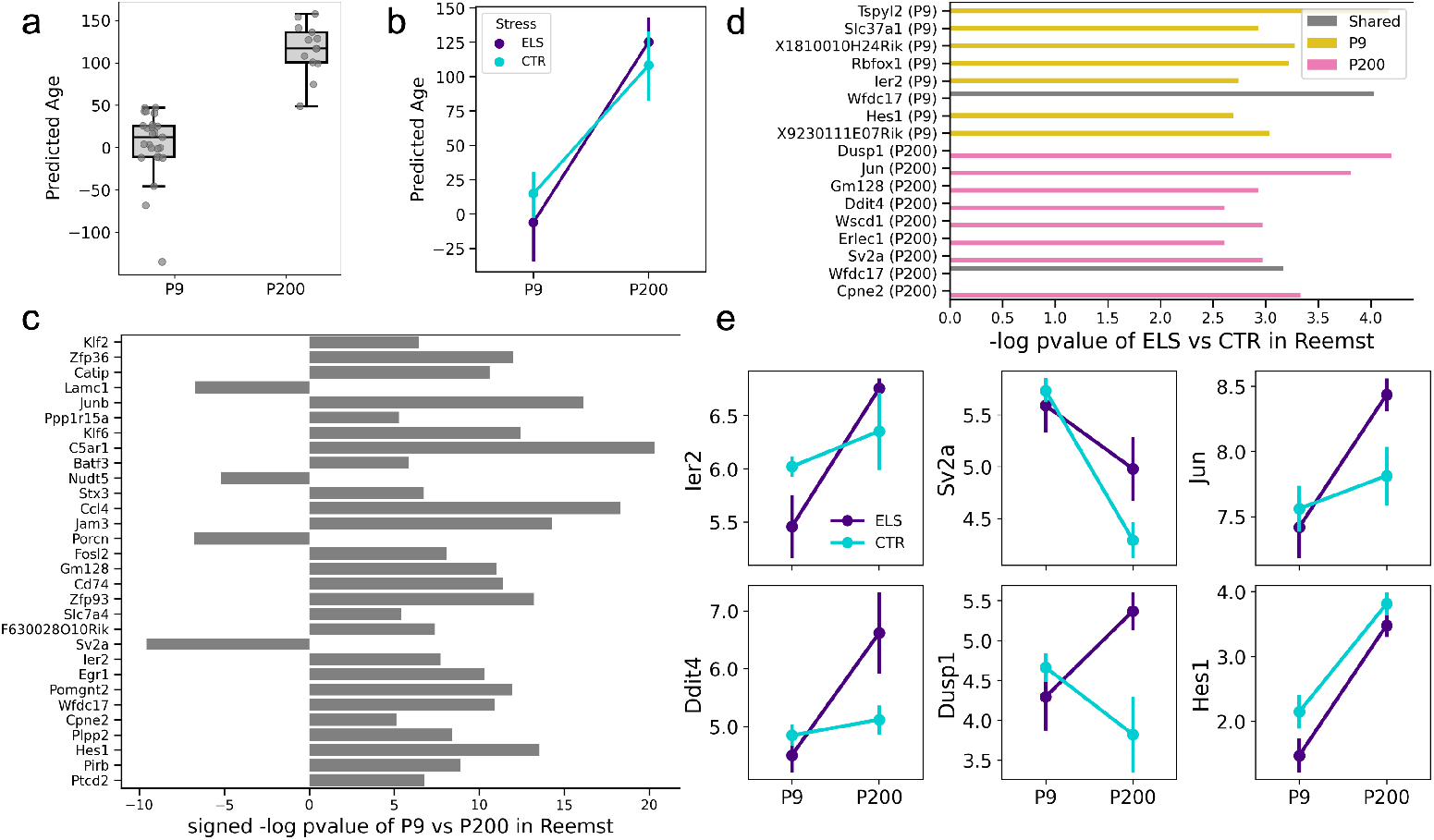
Aging clocks trained on scRNA-seq are applicable to bulk RNA-seq datasets with environmental inputs. **a** Boxplots show the distribution of predicted ages in P9 and P200 mice under the single-cell to bulk-level extrapolation approach. Predicted ages significantly differed between P9 and P200 mice (*p* = 1.69 *×* 10^−10^ under t-test). **b** Samples were separated by stress condition at each age, and line plots visualize the trajectory in mean predicted age (points) at the model between P9 and P200 age groups (linear regression *p* = .19 for P9 and *p* = .37 for P200). Error bars show the standard error of the mean. **c** Barplots visualize the key genes in the elastic net (EN) model trained on the Hammond dataset for predicting age that were also statistically significant (at p-value *<* 10^−5^) for differentiating P9 and P200 mice in the Reemst dataset. Barplots visualize the (signed) −log p-value of each gene under a t-test comparing its expression between P9 and P200 mice in the Reemst dataset. The sign (direction) of the bar indicates the sign of the coefficient of the gene in the EN model for age. **d** Barplots visualize -log p-value (Wilcoxon test) of the ELS vs CTR comparison for the key genes prioritized by the elastic net (EN) model trained on the Hammond dataset. Genes were selected if they had a -log p-value of *>*= 2.5 at ages P9 (gold), P200 (pink) or in both conditions (gray). **e** Line plots visualize expression patterns of key stress-responsive genes, between P9 and P200 mice, separated by ELS (CTR) colored in purple (teal). Highlighted genes with differences between ELS and CTR groups include Ier2 (*p* = 0.064 at P9), Sv2a (*p* = 0.05 at P200), Jun (*p* = 0.02 at P200), Ddit4 (*p* = 0.07 at P200), Dusp1(*p* = 0.015 at P200), and Hes1 (*p* = 0.067 at P9 and *p* = 0.115 at P200)

Frequency was unequivocally the most efficient method, both in terms of run-time and memory requirements, since it is an inherently low-dimensional featurization technique that produces as many features as the number of clusters. The PB++ variants take slightly longer to compute during the featurization step than PBHVG but lead to an overall reduced run-time and memory requirements for the overall pipeline as they produce a low-dimensional space (number of features also equal to the number of clusters). PBHVG produces a large number of features, increasing the run-time and memory requirements. Finally, scLKME has the highest run-time and memory requirements across datasets, likely due to the kernel evaluations built into the method and the high-dimensional feature space produced.

### 2.5 Identifying key dynamic modules of genes changing with the aging trajectory

We next sought to identify the groups of genes that underlie the changes in cell-type frequencies driving the microglia aging clocks. To this end, we used the DELVE algorithm [45] to uncover groups of genes changing dynamically (i.e., increase or decrease in expression) and in characteristic ways during aging. Applying DELVE in the Hammond dataset identified four dynamic gene modules with prominent changes across the aging continuum. As shown in Fig. 6 **left**, key modules and their respective top-scoring genes exhibited characteristic monotonic increases (Fig. 6**a**) or decreases (Fig. 6**b**) in expression over the aging expression over the aging continuum. As a further independent validation, we then examined the expression of representative genes in each module (Fig. 6 **middle**) in an independent bulk RNA-sequencing dataset generated by Reemst *et al*. [42] (hereinafter denoted as the Reemst dataset), profiling microglia from young mice (P9, nine days of age) and older mice (P200, 200 days of age). Strikingly, the expression patterns of genes uncovered in modules in the Hammond dataset tracked similarly in the Reemst dataset (Fig. 6 **middle**), suggesting the trends are robust and generalize between single-cell and bulk data. Similarly, composite expression scores obtained by summing the expression of top five highlighted genes in each module in each mouse are shown in Fig. 6 **right** and show consistent patterns of increase or decrease, according to the DELVE modules.

Analysis with gene ontology run with the g:profiler tool [46] revealed distinct biological processes in each DELVE module. Gene ontology categories enriched in dynamic module 1 included cytoskeleton organization (GO: 0007010), regulation of monocyte differentiation (GO:0045655), and lipopolysaccharide immune receptor activity (GO:0001875) (Supplementary Figure 2). Dynamic module 2 was enriched for generation of precursor metabolites and energy (GO: 0006091), detoxification (GO:0098754), ATP:ADP antiporter activity (GO:0005471), and organonitrogen compound metabolic process (GO:1901564) (Supplementary Figure 3). Dynamic module 3 was enriched for iron-sulfur cluster binding (GO:0051536), mitotic cell cycle (GO:0000278), and ubiquitin protein ligase binding (GO:0031625) (Supplementary Figure 4).

### 2.6 Extrapolating to and validating with bulk RNA-seq datasets

We next tested if microglia aging clocks trained on scRNA-seq data can be applied to independent bulk RNA-seq datasets. As bulk datasets are more readily available, such applicability would present a powerful opportunity to exploit the unique insights gained by single-cell datasets to a wider range of settings. It would further facilitate the use of single-cell-trained markers to dissect how bulk-level gene expression programs relevant to aging are experimentally modulated by various environmental inputs. To address these questions, we leveraged the bulk-RNA seq dataset of microglia generated by Reemst *et al.*. In addition to having different age groups, mice in this study were also exposed to either control (CTR) or early life stress (ELS) in the form of limited bedding.

To assess the bulk-level applicability of scRNA-seq-trained microglia clocks, we extracted the pseudobulk features from the PBHVG aging clock, which was trained earlier in the Hammond dataset, for the 2,845 genes that were also measured in the Reemst dataset. To translate such pseudobulk features into a data structure similar to bulk RNA-seq data with one expression value for each gene per sample, we summed each gene’s per-cluster composite value across all clusters. These per-sample feature vectors of gene expression measurements were then used to train an Elastic Net (EN) regression model for age in the Hammond dataset (see Methods for details). When applied to the Reemst dataset, this single-cell-trained aging clock resulted in predicted ages that differed significantly between P9 and P200 mice (p = 1.69 *×* 10^−10^ under t-test) (Fig. 7**a**). We also tested whether ELS exposure impacted the aging clock. While there were no statistically significant differences between exposure groups (Fig. 7**b**), we noted an intriguing pattern whereby ELS resulted in lower predicted age in P9 (*p* = 0.19) but higher predicted age in P200 as compared to the CTR condition (*p* = 0.37).

As a last step, we examined the expression patterns of key genes prioritized by the PBHVG model across age and stress groups. We specifically considered the subset of genes with a coefficient with an absolute value *>*= 1.2 in the Hammond-trained EN model and a -log p-value *>*= 5 upon testing between P9 and P200 in the Reemst dataset. These analyses identified conserved and distinct transcriptional patterns between P9 and P200 mice, with key genes including *Sv2a,Lamc1,C5ar1, Egr1, Ccl4, Hes1, Batf3, and Zfp93* (Fig. 7**c**). In addition, ELS exposure resulted in distinct gene expression differences between ELS and CTR groups in each age group. P9 mice exposed to ELS showed statistically significant differences in the expression of *Wfdc17* and *Tspyl2* (p *<* 0.05), whereas P200 mice exposed to ELS showed upregulation of *Dusp1* and *Jun* (*p <* 0.05), *Ddit4* (*p* = 0.07), and *Gm128* (*p* = 0.053) (Fig. 7**d**). The dynamic patterns of these stress-driven effects are visualized across the age groups in Fig. 7**e**. Given that no stress-related information was used to train the model, it is intriguing that some of the key genes with expression patterns that change during aging also exhibit expression differences in response to early life stress.

## 3 Discussion

Biological aging clocks have been traditionally developed through bulk-level analyses [3, 3–8], but recent evidence has highlighted the importance of gaining cell-type and single-cell-level insights into the aging process [9–14]. Interrogating the transcriptome of key immune cell types, such as microglia [15, 16, 19, 23, 26], may hold particular promise for gaining such insights. Leveraging scRNA-seq datasets and state-of-the-art computational methods, we generated robust microglia-derived aging markers, compared marker performance across different featurization approaches, identified microglia genetic programs that change dynamically with aging, and showed that single-cell insights extrapolate to bulk RNA-seq data.

Rigorous testing of classification and regression-based aging markers in the three single-cell datasets highlighted ways in which signals across individual cells, samples, and datasets can be integrated to generate machine learning models of dynamic processes such as aging clocks. Leveraging a variety of such approaches, including frequency [32, 33], scLKME [39], our novel pseudobulk++ approach, and classical pseudobulk [40], revealed intriguing tradeoffs between accuracy, biological interpretability, and efficiency. Frequency and pseudobulk++ compute compact, information-rich features per each microglia subtype, capturing the extent to which their abundances and gene expression patterns correlate with age. Frequency and pseudobulk++ proved to be top performers in age classification in the Hammond and Kracht datasets, and among the top two performing methods in regression-based aging clocks in the Buckley and Hammond datasets that have samples from donors across a wide age range. In contrast, scLKME and classical pseudobulk produce higher-dimensional data representations, which inevitably pose larger demands on computation time and memory, but may encode more nuanced information that improves prediction in particular settings. For example, in the Buckley dataset, scLKME had significantly higher accuracy as an age classifier than the other methods, suggesting that the higher dimensional feature space was advantageous in this dataset. Similarly, PBHVG and scLKME may be more accurate as aging clocks in challenging prediction tasks with limited age ranges, such as in the Kracht dataset. As our study only examines unsupervised approaches, there are promising opportunities to further explore how supervised, learning-based featurization [36, 37] and their variants incorporating additional donor-level covariates [38] can be applied to create more information-rich encodings of single-cell profiles.

Prior work has supported central roles for microglia in the neuroinflammation associated with aging [17, 18], neurodegenerative states [21–23], and diverse neuropsychiatric diseases [24, 25]. However, the specific microglia subtypes and biological processes underlying these roles are unclear. Our unsupervised machine learning analyses, applied across three independent datasets, indicate dynamic age-related changes in the frequency of microglia subtypes, which are marked by biologically relevant key signatures. For example, *Apoe* encodes an apolipoprotein that has a well established role in neurodegeneration and dementia [47], and *Malat1* codes for a long non-coding RNA that is dysregulated in immune cell subtypes in association with aging and frailty [48]. These findings corroborate and extend recent scRNA-seq studies showing that transcriptional alterations during aging involve specific microglia subtypes and key genetic programs relevant to aging-related brain phenotypes, including neural stem cell proliferation, neurodegeneration, and Alzheimer’s disease [27–31]. Applying dynamic gene module (DELVE [45]) and ontology analyses further identified gene modules enriched for distinct biological processes, including cytoskeleton organization, immune cell differentiation, metabolism-related processes, cell cycle regulation, and ubiquitination. Together these findings suggest promising research avenues to uncover the role of microglia in brain aging and disease.

Our results also indicate that single-cell-trained aging clocks are applicable to bulk-level data. More specifically, we found that our pseudobulk clock trained from scRNA-seq robustly predicted age in an independent bulk RNA-seq dataset. Consistent with prior work showing convergent genomic effects of stress and aging [49–51], we further found that ELS significantly influenced the bulk-level expression of several top genes comprising the pseudobulk clock, with effect magnitudes differing between age groups. Notable examples include the genes encoding the master transcription factor AP-1 (*Jun*) [52] and the key innate immunity regulator *Dusp1* [53]. This suggests that single-cell-derived transcriptomic signatures of aging are applicable to bulk data and modulated by environmental input. If confirmed by future studies, such single-cell to bulk-level applicability presents powerful opportunities to exploit the unique biological insights gained by single-cell datasets to a wider range of settings, spanning large-scale human cohorts and more nuanced experimental systems.

In summary, the present study builds on the highly promising biological aging research by leveraging scRNA-seq data and state-of-the-art machine learning methods to identify robust microglia aging clocks and dynamic cell-type-specific genetic programs. Such single-cell-derived and cell-type-specific clocks can yield unique insights into brain aging, ultimately promoting interventions that beneficially modulate health and disease trajectories.

## 4 Methods

### 4.1 Data Acquisition and Pre-Processing

#### Single-Cell Datasets

The Hammond [27] (GSE121654), Buckley [28] (GSE196364), and Kracht [41] (GSE141862) scRNA-seq datasets were downloaded from Gene Expression omnibus (GEO). Associated accession numbers are indicated in parentheses. Note that in the Buckley dataset, we converted the Seurat `multi_intergrated_seurat2020.rds` object available in Zenodo (https://zenodo.org/records/7145399) as mentioned in Ref. [28] to an AnnData object for our experiments.

We performed downsampling in each dataset to increase computational efficiency. In each dataset, for a given sample *i*, max{750, total number of cells in sample i} were randomly selected. Cells across samples were then concatenated into an `annData` object for use with single-cell pre-processing tools in `Scanpy`. In each sample, we performed Counts per million (CPM) normalization by normalizing each cell by the total number of counts across all genes, using the function `scanpy.pp.normalize_total` with a target sum of 1e6. We then performed log(1 + *x*) transformation for all counts. In the Hammond and Buckley datasets, the 3,000 most highly variable genes were retained. In the Kracht dataset, there were only 1,685 measured genes, and hence we did not do any additional highly variable gene filtering.

#### Reemst Bulk RNA Sequencing Dataset

The Reemst bulk RNA-sequencing dataset [42] was downloaded from GEO (accession number GSE207067). All gene expression measurements were log(1 + *x*) transformed. For all experiments in the paper, we did not consider any sample from a mouse that had received a Lipopolysaccharide (LPS) injection.

### 4.2 Notation and Preliminaries

We define an *N* profiled sample single-cell dataset, *X* of *N* cell *×* gene matrices as 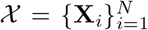 Here, 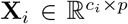 gives the data matrix of *p* transcriptomic features measured across *c*_*i*_ cells in a particular sample, *i*. We furthermore define a vector of per-sample ages, **y** *∈* ℝ^*N*^, such that the *i*-th element, *y*_*i*_ gives the age of profiled sample *i*. Moreover, our task is to employ a robust featurization strategy to create a per-sample microglia transcriptomic summary given by *d* features, **s**_*i*_ ∈ ℝ^*d*^, such that some model, *f* (*·*) can accurately model age so that 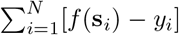 is as small as possible.

#### Defining Cell Types Through Unsupervised Clustering

The partitioning of cells into clusters forms the backbone of many of the featurization approaches, which engineer sample-level features based on clusters or uncovered cell types. To pre-process cells for input to clustering algorithms, we used `scanpy` to first represent each cell in terms of its top 40 principal components (`PCs = 40`) and built a *k*-nearest neighbor graph (`kNN=10`) by connecting each cell to its 10 nearest neighbors. To partition single cells into salient cell types with common gene expression programs, we used Leiden clustering [54] on constructed graph representation of the data, with resolution parameter, *γ* = 1. This algorithm partitions cells in each sample **X**_*i*_ into *K* populations. Note that this default resolution parameter, *γ* will produce different numbers of clusters per dataset, depending on the total number of cells and the extent of heterogeneity across cells.

### 4.3 Featurization Strategies

#### Frequency

Given a partitioning of cells in a given sample, **X**_*i*_ into 1 of *K* populations, we engineer a frequency feature vector, **f** ^*i*^, for sample *i*, such that the *m*-th entry, 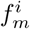 encodes the proportion of cells in **X**_*i*_ that were assigned to cluster *m*. This implies that the frequencies sum to 1 with 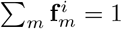

#### Classical Pseudobulk (PBHVG)

Classical psuedobulk is a standard formulation for compressing aggregate information across cells into single per-sample feature vectors. Here, we compute classical pseudobulk based only on the most highly variable genes in each dataset, such that for a given sample, **X**_*i*_, the objective is to look at summed expressions of each gene across all cells within a particular cluster, *m*. This process is repeated across the *K* clusters. So, we can define a pseudobulk feature vector for sample, *i* as **p**^*i*^ by concatenating their per-each-cluster feature representations (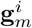 for cluster *m* in sample *i*) as,

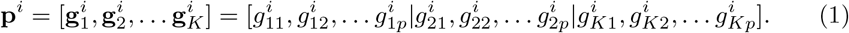

Here, for a dataset with *p* genes, each 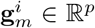 is a vector of aggregated expressions over the *p* measured genes from all cells in sample *i* that have been assigned to cluster *m*. We chose sum as our aggregation technique so that a particular value, 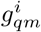 gives the sum of expression of gene *q* of all cells in cluster *m* in sample *i*.

#### Pseudobulk++

Pseudobulk++ (PB++) is a novel featurization strategy introduced here, which is a parsimonious formulation of combining notions of frequency with classical pseudobulk. For each sample, this method computes a score for each cluster reflecting both the frequency of the sample’s cells for that cluster and the extent to which the sample’s cells in that cluster express key genes.

First, we perform a differential expression test in each cluster, which implements a *cluster vs. rest* comparison in each of the *K* clusters. We then rank genes by p-value from most to least significant, such that the genes at the top of the list are those that are significantly differentially expressed through prominent up-regulation or down-regulation in at least one cluster. We then extract the top *G* genes, in terms of their p-values (and z-scores) across clusters to form the vector, **f**, such that **f** = [*f*_1_, …, *f*_*G*_] *∈* ℝ^*G*^. Here, *f*_*q*_ gives the absolute value of the z-score for gene *q*, reflecting its strength of differential expression (either significantly up or down regulated) in the one vs. rest differential expression test in at least one the clusters. To implement differential expression tests, we used `get.rank_genes_groups` in `scanpy`. In practice, we found the method works well for the top 25 or top 50 differentially expressed genes and their respective scores across all clusters to yield *G* = 25 and *G* = 50 for PB++25 and PB++50, respectively. Ultimately, the set of *G* genes will be treated as a common set of key genes used to compute a composite per-cluster score for each sample.

For a particular sample, *i*, we extract a subset of their cell *×* gene matrix from **X**_*i*_ as 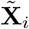 with 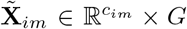, which gives the expression of only the common top *G* genes across the *c*_*im*_ cells in cluster *m* in sample *i*. The feature value, 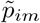 for cluster *m* in sample *i* is therefore computed as,

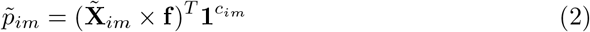

Here, 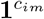 is notation for the vector of length *c*_*im*_ (e.g. the number of cells from sample *i* in cluster *m* and provides an operator to sum the re-weighted values of cells attained by 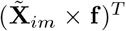. Moreover, the entire feature vector for sample *i* under the PB++ formulation can be expressed as,

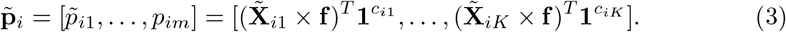

Ultimately, each 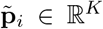 is relatively low dimensional and only has as many features as there are clusters.

#### scLKME

The scLKME strategy was introduced in Ref. [39]. The premise of the algorithm is to choose a number of landmark cells across all profiled samples and to ultimately compute featurizations based on overall patterns of each cell’s similarity with each landmark as gleaned through kernel evaluations. We chose *L* = 1500 landmark cells obtained through cell sketching [55] across all samples and with kernel, *κ*(*·*), as the radial basis function (RBF). Under this formulation, scLKME computes the featurization for sample *i*, **s**_*i*_ *∈* ℝ^*L*^ (for *L* = 1500) as,

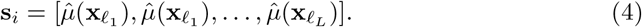

Here, each 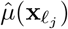 computes the mean kernel evaluation over all *c*_*i*_ cells and sample *i* and the landmark cell, *l*_*j*_ as,

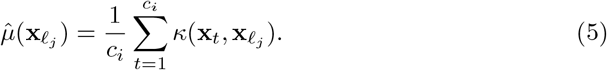

### 4.4 Age classifiers

Age classifiers were trained to predict the ages of samples binned in discrete categories. The Hammond dataset already had discrete age categories, consisting of E14.5, P4, P5, P30, P100, and P540, so the classification problem was over six classes. In the Buckley dataset, we binned ages into young (*<* 4 months), adult (4 months ≤ age *<* 14), and old (≥ 14 months). The classification problem was therefore over three age categories. Finally, we formulated a binary classification problem in the Kracht dataset by separating samples into first (9-12 weeks) and second trimester (13-18 weeks), respectively.

Within a dataset, we applied a given featurization approach to each sample to obtain their vector encoding. These vector encodings were then given to a random forest classifier (implemented in `ScikitLearn` using `RandomForestClassifier` in Python) with 50 trees and the square root of the total number of features to find the best split. We used 200 randomized train/test splits of the data using 80% of the donors and their respective samples (in the Hammond and Kracht dataset, which have multiple samples per donor) or samples (in the Buckley dataset) for training and the remaining 20% for testing to obtain a distribution of classification accuracies. Classification accuracies reflect the proportion of samples with correctly predicted labels in the dataset.

### 4.5 Aging clocks

We used penalized linear regression with the Lasso penalty (implemented in `ScikitLearn` using `linear_model.Lasso` in Python) to predict sample ages, under each given featurization. Given a featurization of *N* samples into *d* features obtained for a dataset that results in a matrix, **D** *∈* ℝ^*N×f*^ with a corresponding age response vector as **y** *∈* ℝ^*N*^, Lasso regression seeks to optimize a vector of per-feature coefficients, *β ∈* ℝ^*f*^ that minimize,

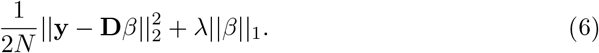

We tuned *λ*, or the magnitude of the penalty applied to each coefficient, through cross-validation on each training set. To train and test the models, we used a leave-one-out cross-validation approach (LOOCV) by training the model on all but the one held-out sample and predicting the age for the held-out sample in each LOOCV iteration. The Buckley dataset only has one sample per donor, whereas the Kracht and Hammond datasets have multiple samples per donor, such that samples from a donor were kept together in the same train/test split in each LOOCV iteration.

We used Pearson correlation and fitted linear regression p-value as the metrics of success to quantify how well the chronological and predicted ages correlated across samples (implemented with `scikit learn`).

### 4.6 Extrapolating from Single-Cell to Bulk

To generate a scRNA-seq-derived aging clock that could be applied to the Reemst bulk RNA-seq dataset [42], we converted the multiple cell *×* gene expression matrices in the single-cell Hammond dataset [27] into a sample *×* gene expression matrix, which has analogous structure to the bulk RNA-seq data. To this end, we used a simple variation of the classical pseudobulk approach. As classical pseudobulk computes an aggregate expression measurement for each gene in each cluster (usually by computing a sum), we decided to simply compute aggregate expression of each gene across all cells (so, across all clusters) through sum (alternative aggregation methods explored in Supplementary Figure 5). So, given the Hammond dataset, 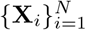 profiling cells across *N*_*h*_ samples with *c*_*i*_ cells measured per sample, we computed the sample *×* gene matrix, 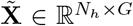 for the dataset such that a given entry 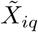 measuring the expression of gene *q* in sample *i* is computed as the sum of column *q* (gene *q*) over all cells,

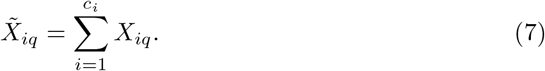

We consider only the *G* genes that are also measured in the Reemst Bulk RNA-sequencing dataset profiling *N*_*r*_ samples, which produced a data matrix of 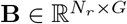.

Using 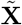 we trained an Elastic Net linear regression model to find per-gene coefficients *β ∈* ℝ^*G*^ to optimally predict ages in the Hammond dataset given by 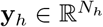 that minimize,

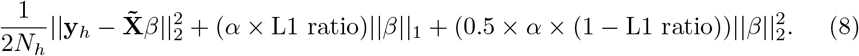

The model is trained using `linear_model.ElasticNet` in `Scikit learn` with default parameter values of `alpha = 1` and `l1_ratio=0.5`.

The ultimate per-gene coefficients encoded in *β* were ultimately used to predict the ages 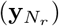 of samples in the Reemst dataset as,

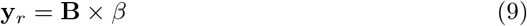

### 4.7 DELVE implementation to identify gene modules with common genetic patterns

We applied the `DELVE` algorithm [45] to the Hammond dataset to reveal dynamic genetic programs in microglia in an unsupervised manner. We uncovered 5 dynamic modules with DELVE (using parameter `n_clusters=5`) and otherwise default parameters.

## 5 Data and Code Availability

All featurization strategies and tutorials for reproducing results are available in github https://github.com/CompCy-lab/microglia-aging-clock. Processed scRNA-seq datasets are available in anndata format in Zenodo (DOI: 10.5281/zenodo.12811383).

## 6 Author Contributions

N.S. and A.Z. conceptualized the study and designed experiments. Featurization techniques were developed and implemented by N.S., A.Z., L.D., and Y.H. Datasets were pre-processed by L.D., Y.H., and S.J. Data were analyzed by all authors. N.S., A.Z., and L.D. wrote the paper with input from all authors.

## 7 Acknowledgements

N.S. acknowledges support from the National Institute on Aging of the National Institute of Health (NIH) under award 1R21AG084251-01A1.

