## Supplementary Information for "Leveraging Single-Cell RNA-Seq to Generate Robust Microglia Aging Clocks"

### Supplementary Material

November 6, 2024

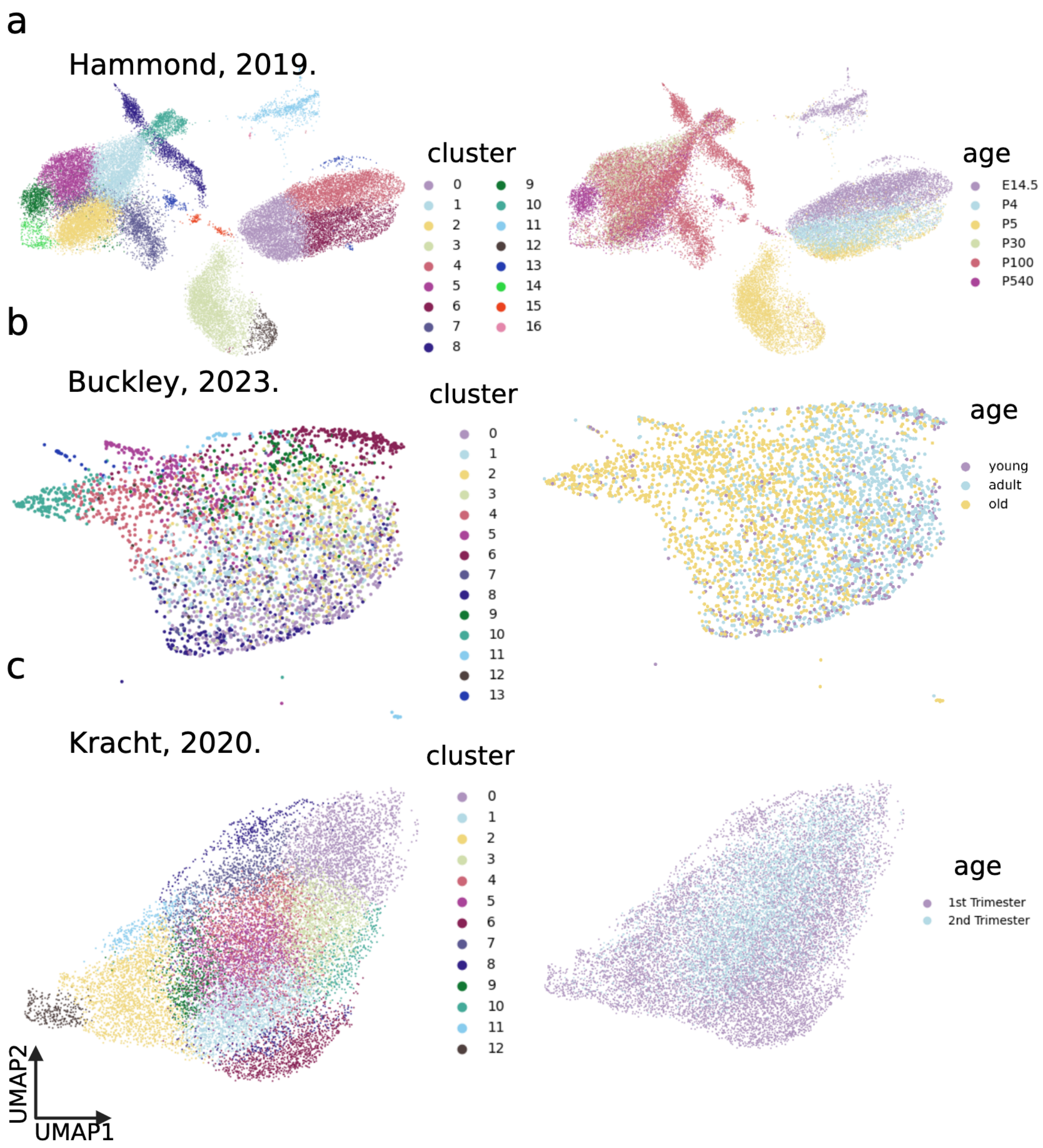


Figure 1: **Clustering and two-dimensional visualization of cells across datasets reveal microglia subtypes with characteristic genetic programs**. Cells were projected in two dimensions with UMAP and colored according to Leiden cluster (**left**) and age group (**right**) in the Hammond **a**, Buckley **b**, and Kracht **c** datasets.

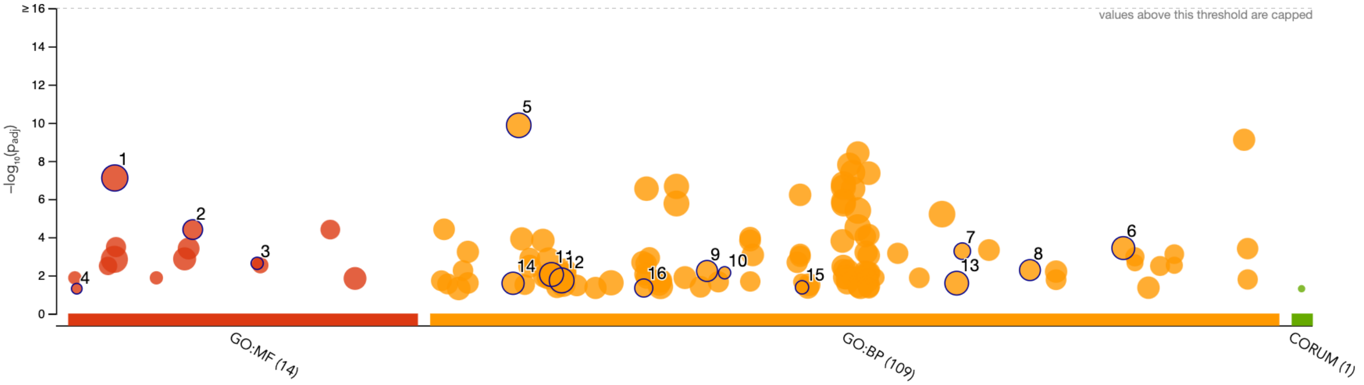


| ID | Source | Term ID | Term Name | $p_{adj}$ (query_1) |
|---|---|---|---|---|
| 1 | GO:MF | GO:0005515 | protein binding | $7.829\times10^{-8}$ |
| 2 | GO:MF | GO:0030695 | GTPase regulator activity | $3.968\times10^{-5}$ |
| 3 | GO:MF | GO:0045028 | G protein-coupled purinergic nucleotide recept... | $2.330\times10^{-3}$ |
| 4 | GO:MF | GO:0001875 | lipopolysaccharide immune receptor activity | $4.971\times10^{-2}$ |
| 5 | GO:BP | GO:0007275 | multicellular organism development | $1.355\times10^{-10}$ |
| 6 | GO:BP | GO:1901700 | response to oxygen-containing compound | $3.695\times10^{-4}$ |
| 7 | GO:BP | GO:0071277 | cellular response to calcium ion | $5.385\times10^{-4}$ |
| 8 | GO:BP | GO:0097435 | supramolecular fiber organization | $5.283\times10^{-3}$ |
| 9 | GO:BP | GO:0034330 | cell junction organization | $5.848\times10^{-3}$ |
| 10 | GO:BP | GO:0035589 | G protein-coupled purinergic nucleotide recept... | $7.294\times10^{-3}$ |
| 11 | GO:BP | GO:0010033 | response to organic substance | $9.026\times10^{-3}$ |
| 12 | GO:BP | GO:0010468 | regulation of gene expression | $1.834\times10^{-2}$ |
| 13 | GO:BP | GO:0070887 | cellular response to chemical stimulus | $2.559\times10^{-2}$ |
| 14 | GO:BP | GO:0007010 | cytoskeleton organization | $2.588\times10^{-2}$ |
| 15 | GO:BP | GO:0045655 | regulation of monocyte differentiation | $4.214\times10^{-2}$ |
| 16 | GO:BP | GO:0022408 | negative regulation of cell-cell adhesion | $4.588\times10^{-2}$ |

Figure 2: **Gene ontology results for genes in module 1 under DELVE.** Top enriched GO-terms among genes in dynamic module 1 (increasing with age) obtained with DELVE in the Hammond dataset.

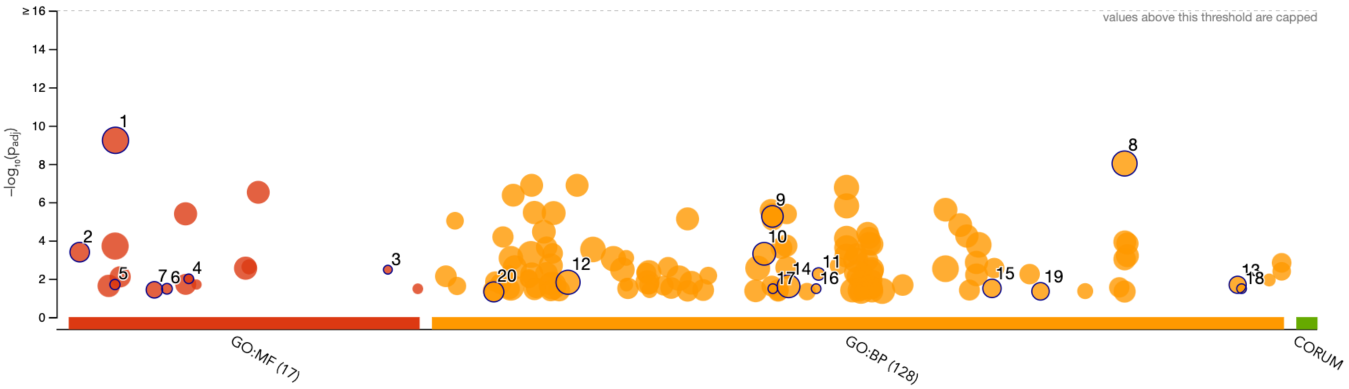


| ID | Source | Term ID | Term Name | $p_{adj}$ (query_1) |
|---|---|---|---|---|
| 1 | GO:MF | GO:0005515 | protein binding | $5.872\times10^{-10}$ |
| 2 | GO:MF | GO:0003729 | mRNA binding | $4.058\times10^{-4}$ |
| 3 | GO:MF | GO:0106130 | purine phosphoribosyltransferase activity | $3.345\times10^{-3}$ |
| 4 | GO:MF | GO:0030060 | L-malate dehydrogenase activity | $1.002\times10^{-2}$ |
| 5 | GO:MF | GO:0005471 | ATP:ADP antiporter activity | $2.000\times10^{-2}$ |
| 6 | GO:MF | GO:0017077 | oxidative phosphorylation uncoupler activity | $3.328\times10^{-2}$ |
| 7 | GO:MF | GO:0016209 | antioxidant activity | $3.766\times10^{-2}$ |
| 8 | GO:BP | GO:1901564 | organonitrogen compound metabolic process | $9.509\times10^{-9}$ |
| 9 | GO:BP | GO:0043066 | negative regulation of apoptotic process | $5.510\times10^{-6}$ |
| 10 | GO:BP | GO:0042592 | homeostatic process | $4.956\times10^{-4}$ |
| 11 | GO:BP | GO:0046166 | glyceraldehyde-3-phosphate biosynthetic pro... | $5.629\times10^{-3}$ |
| 12 | GO:BP | GO:0010646 | regulation of cell communication | $1.519\times10^{-2}$ |
| 13 | GO:BP | GO:1905954 | positive regulation of lipid localization | $2.061\times10^{-2}$ |
| 14 | GO:BP | GO:0044419 | biological process involved in interspecies inte... | $2.455\times10^{-2}$ |
| 15 | GO:BP | GO:0072330 | monocarboxylic acid biosynthetic process | $3.175\times10^{-2}$ |
| 16 | GO:BP | GO:0046083 | adenine metabolic process | $3.278\times10^{-2}$ |
| 17 | GO:BP | GO:0043096 | purine nucleobase salvage | $3.278\times10^{-2}$ |
| 18 | GO:BP | GO:1990428 | miRNA transport | $3.278\times10^{-2}$ |
| 19 | GO:BP | GO:0098754 | detoxification | $4.522\times10^{-2}$ |
| 20 | GO:BP | GO:0006091 | generation of precursor metabolites and energy | $4.838\times10^{-2}$ |

Figure 3: **Gene ontology results for genes in module 2 under DELVE.** Top enriched GO-terms among genes in dynamic module 2 (decreasing with age) obtained with DELVE in the Hammond dataset.

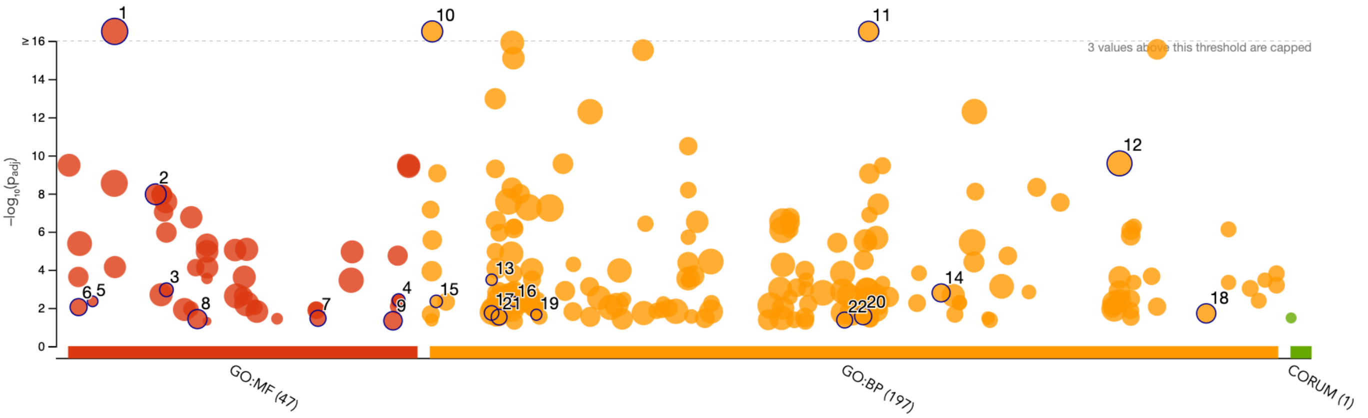


| ID | Source | Term ID | Term Name | $p_{adj}$ (query_1) |
|---|---|---|---|---|
| 1 | GO:MF | GO:0005515 | protein binding | $4.243\times10^{-18}$ |
| 2 | GO:MF | GO:0016462 | pyrophosphatase activity | $1.092\times10^{-8}$ |
| 3 | GO:MF | GO:0017116 | single-stranded DNA helicase activity | $1.111\times10^{-3}$ |
| 4 | GO:MF | GO:0140713 | histone chaperone activity | $3.871\times10^{-3}$ |
| 5 | GO:MF | GO:0004448 | isocitrate dehydrogenase [NAD(P)+] activity | $4.490\times10^{-3}$ |
| 6 | GO:MF | GO:0003697 | single-stranded DNA binding | $9.014\times10^{-3}$ |
| 7 | GO:MF | GO:0051536 | iron-sulfur cluster binding | $3.478\times10^{-2}$ |
| 8 | GO:MF | GO:0031625 | ubiquitin protein ligase binding | $3.829\times10^{-2}$ |
| 9 | GO:MF | GO:0140097 | catalytic activity, acting on DNA | $4.675\times10^{-2}$ |
| 10 | GO:BP | GO:0000278 | mitotic cell cycle | $5.016\times10^{-18}$ |
| 11 | GO:BP | GO:0051276 | chromosome organization | $2.416\times10^{-17}$ |
| 12 | GO:BP | GO:1901564 | organonitrogen compound metabolic process | $2.626\times10^{-10}$ |
| 13 | GO:BP | GO:0006102 | isocitrate metabolic process | $3.295\times10^{-4}$ |
| 14 | GO:BP | GO:0065004 | protein-DNA complex assembly | $1.621\times10^{-3}$ |
| 15 | GO:BP | GO:0000727 | double-strand break repair via break-induced r... | $4.507\times10^{-3}$ |
| 16 | GO:BP | GO:0007057 | spindle assembly involved in female meiosis I | $4.572\times10^{-3}$ |
| 17 | GO:BP | GO:0006099 | tricarboxylic acid cycle | $1.819\times10^{-2}$ |
| 18 | GO:BP | GO:1904951 | positive regulation of establishment of protein l... | $1.924\times10^{-2}$ |
| 19 | GO:BP | GO:0009211 | pyrimidine deoxyribonucleoside triphosphate ... | $2.257\times10^{-2}$ |
| 20 | GO:BP | GO:0051028 | mRNA transport | $2.674\times10^{-2}$ |
| 21 | GO:BP | GO:0006446 | regulation of translational initiation | $3.003\times10^{-2}$ |
| 22 | GO:BP | GO:0048599 | oocyte development | $4.288\times10^{-2}$ |

Figure 4: **Gene ontology results for genes in module 3 under DELVE.** Top enriched GO-terms among genes in dynamic module 3 (marginally decreasing with age) obtained with DELVE in the Hammond dataset.

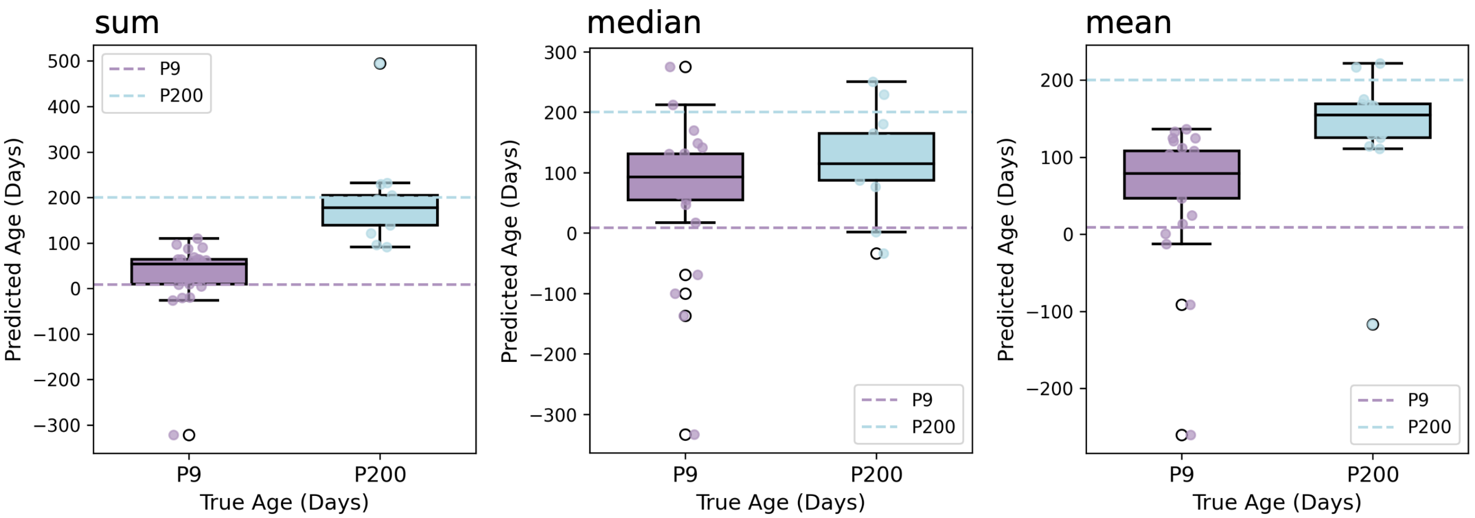


Figure 5: **Possible pooling methods for generating simplified pseudobulk.** Various pooling approaches, including sum (**left**), median (**center**), and mean (**right**) were explored for converting single-cell profiles into a simplified pseudobulk data structure. Briefly, these approaches indicate how the expression of genes are aggregated across cells in samples profiled with single-cell to create a sample $\times$ gene data structure that can be used to train a model, which can be applied to a bulk RNA-seq dataset.

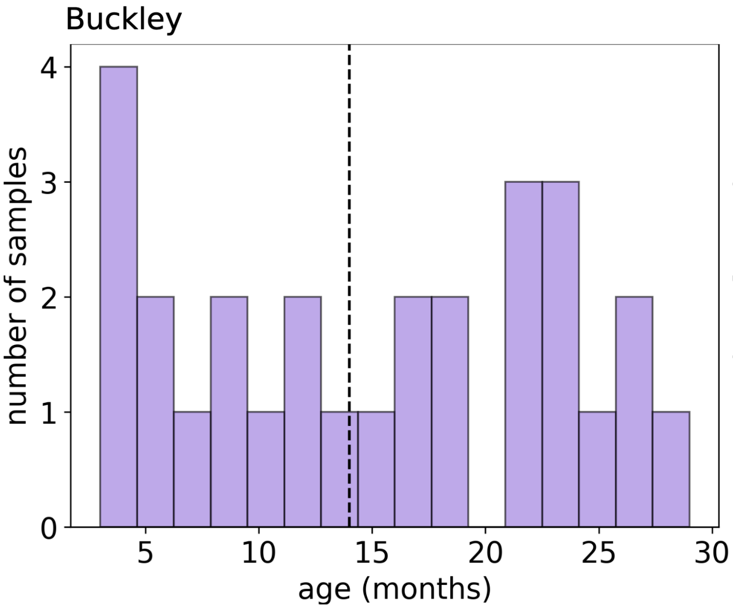


Figure 6: **Distribution of age in the Buckley and Kracht datasets.** To formulate age classification problems, ages in the Buckley and Kracht datasets were binned into two discrete age categories. Histograms show the distribution of edges across samples and the dotted vertical line shows the age threshold used to bin samples by age. Note that samples were split by trimester (first vs second) in the Kracht dataset.

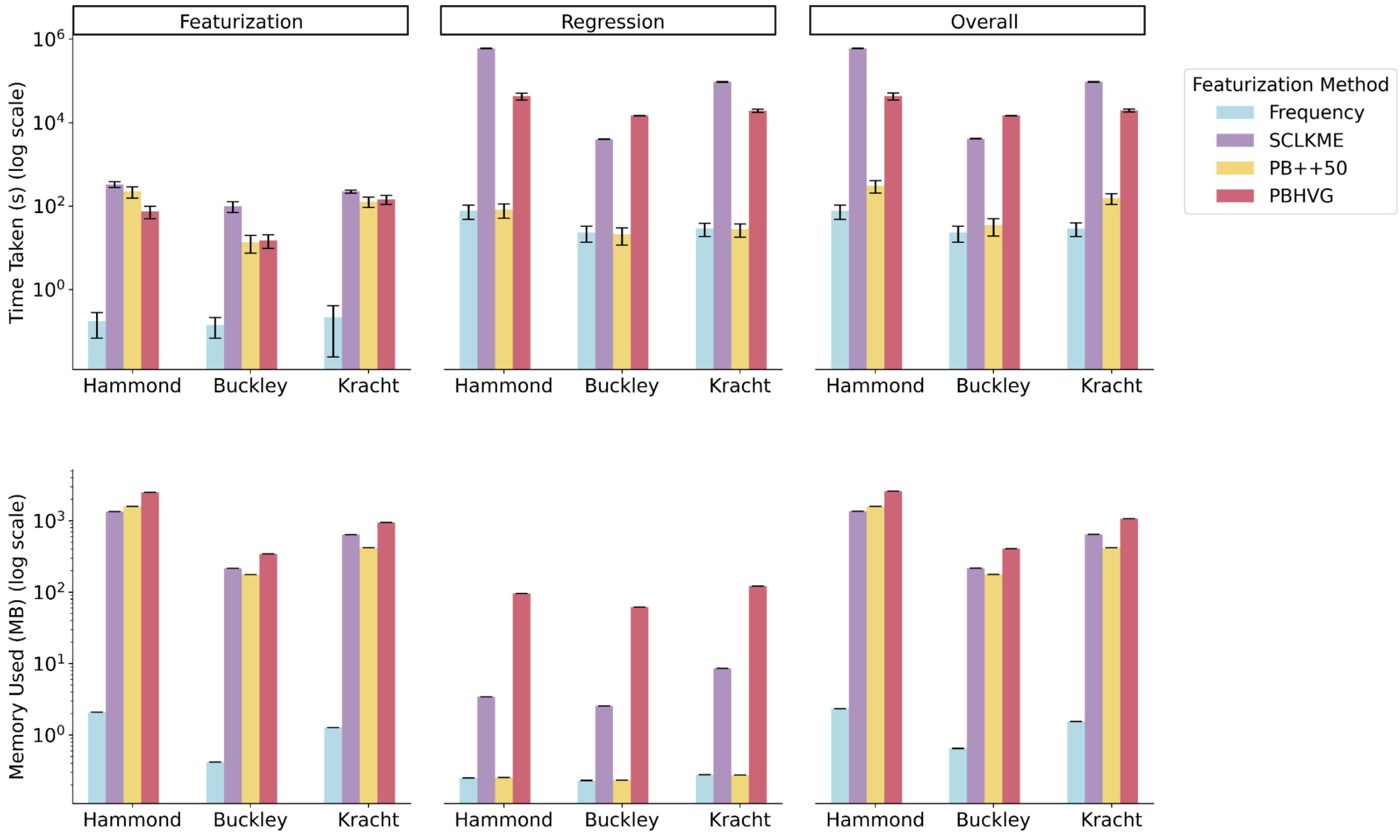


Figure 7: **Run-time and memory requirements for regression-based aging clocks.** Run-time (**top**) and memory (**bottom**) required for each featurization approach, including frequency, scLKME, pseudobulk++50 (PB++50), and pseudobulk with highly variable genes (PBHVG) were evaluated across datasets for featurization only (**left**), Lasso regression to fit the clock (**middle**), and the overall process of featurization and regression (**right**). Results were obtained by repeating the featurization and model training over 30 trials and reporting the mean. Error bars show standard deviation around the mean.

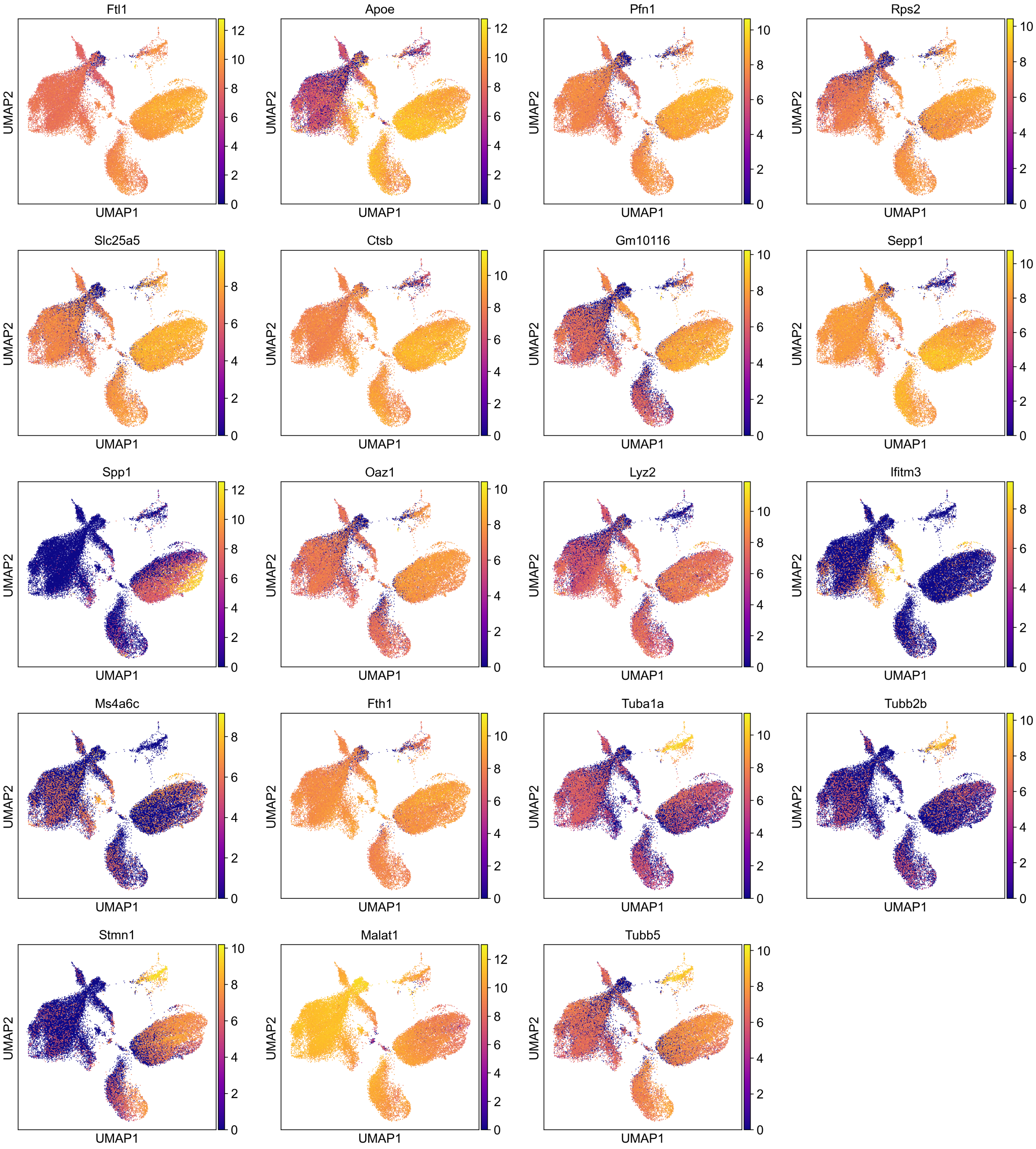


Figure 8: **Key differentially expressed genes across clusters in the Hammond dataset.** Cells from all samples in the Hammond dataset were projected in two dimensions with UMAP and colored by the expression of key cluster-specific differentially expressed genes.

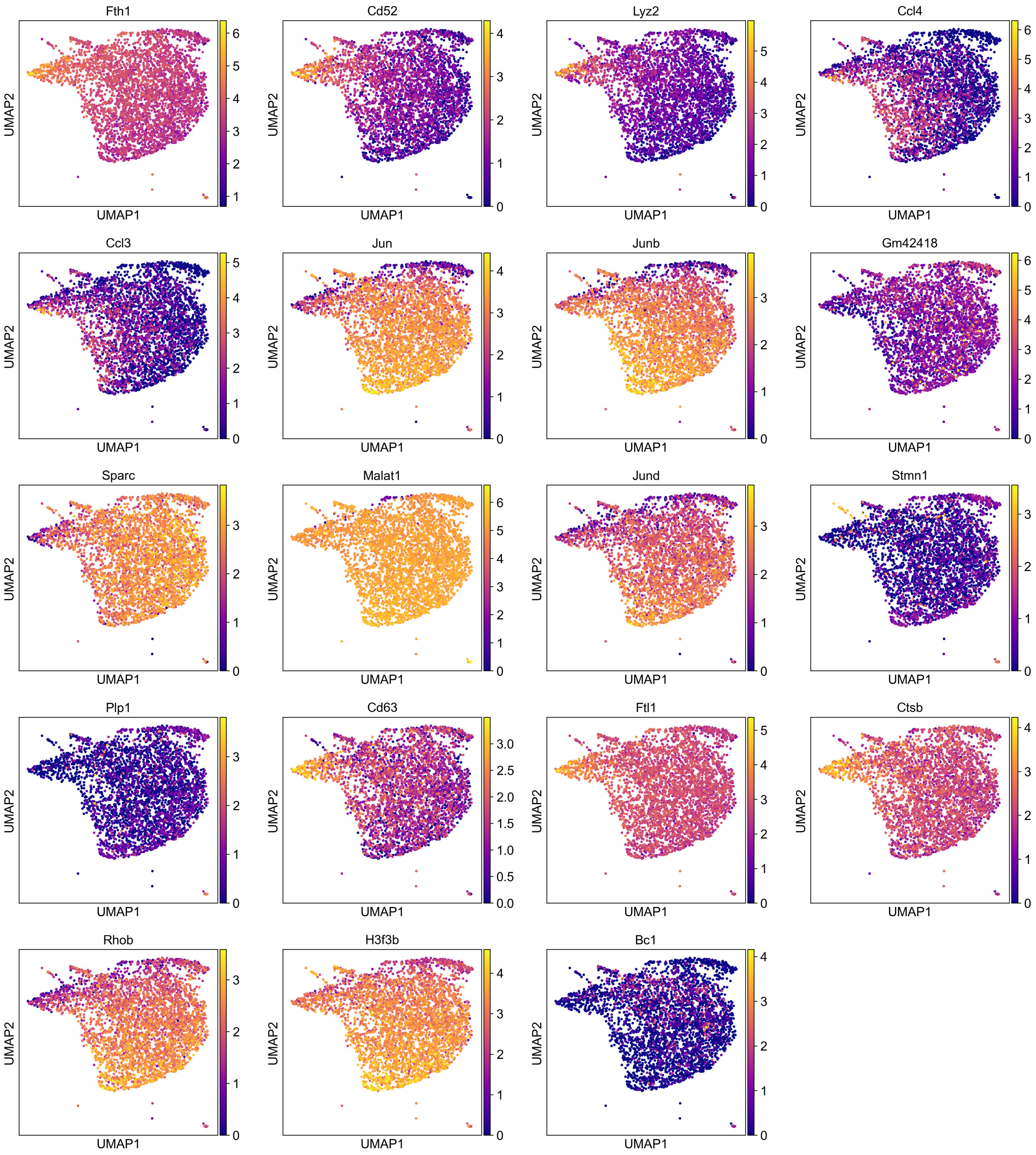


Figure 9: **Key differentially expressed genes across clusters in the Buckley dataset.** Cells from all samples in the Buckley dataset were projected in two dimensions with UMAP and colored by the expression of key cluster-specific differentially expressed genes.

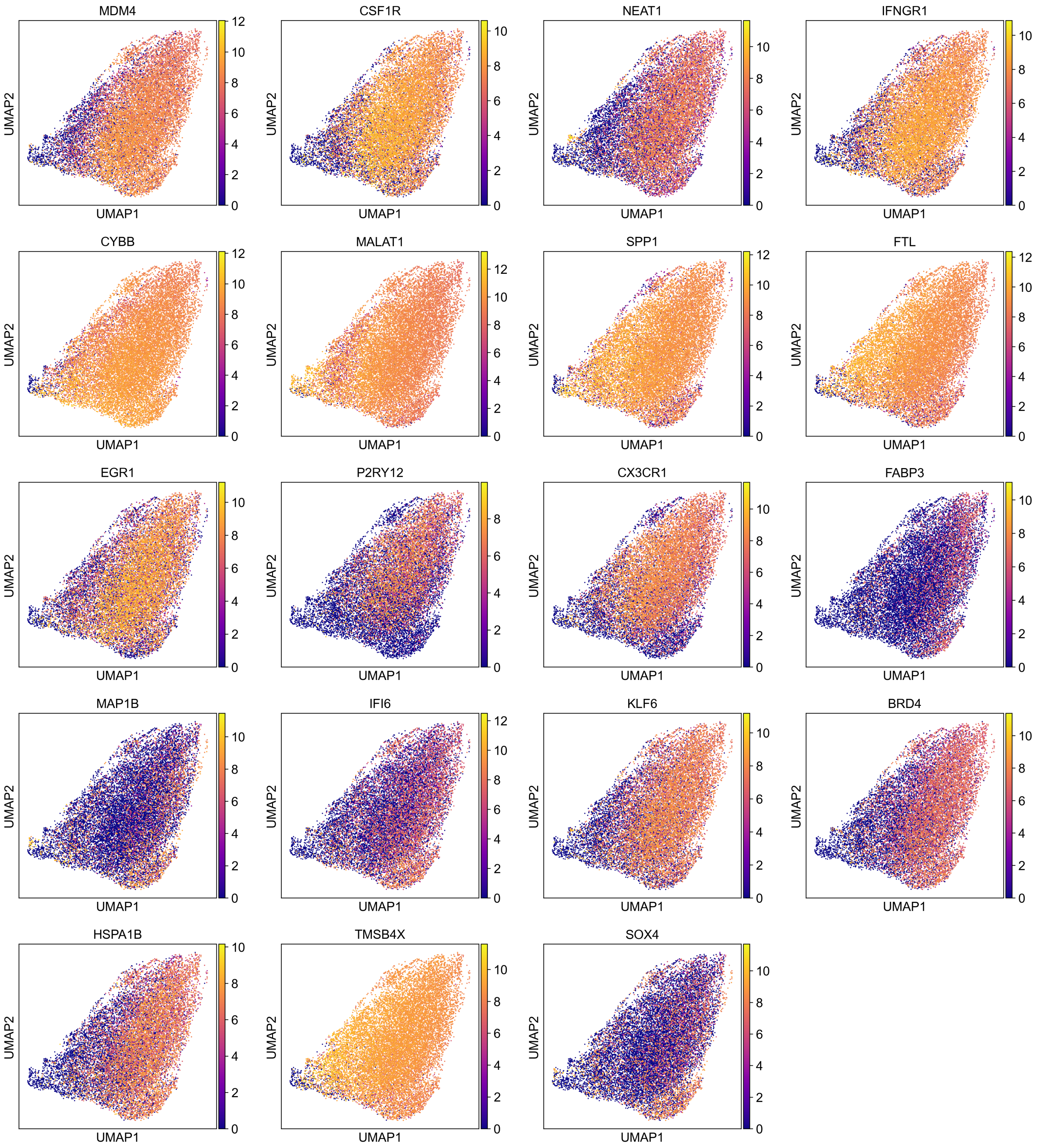


Figure 10: **Key differentially expressed genes across clusters in the Kracht dataset.** Cells from all samples in the Kracht dataset were projected in two dimensions with UMAP and colored by the expression of key cluster-specific differentially expressed genes.
